# Cryo-EM structure of cell-free synthesized human histamine H_2_ receptor coupled to heterotrimeric G_s_ protein in lipid nanodisc environment

**DOI:** 10.1101/2023.07.27.550782

**Authors:** Zoe Köck, Kilian Schnelle, Margherita Persechino, Simon Umbach, Hannes Schihada, Dovile Januliene, Kristian Parey, Steffen Pockes, Peter Kolb, Volker Dötsch, Arne Möller, Daniel Hilger, Frank Bernhard

**Author notes:** authors contributed equally (Zoe Köck, Kilian Schnelle). Correspondence: Arne Möller, Daniel Hilger, Frank Bernhard. **Conflict of interest:** The authors declare that the research was conducted in the absence of any commercial or financial relationships that could be construed as a potential conflict of interest.

## Abstract

Here we describe the cryo-electron microscopy structure of the human histamine 2 receptor (H_2_R) in an active conformation with bound histamine and in complex with G_s_ heterotrimeric protein at an overall resolution of 3.4 Å. The complex was generated by cotranslational insertion into preformed nanodisc membranes using cell-free synthesis in *E. coli* lysates. It is the first structure obtained by this detergent-free strategy and the first GPCR/G_s_ complex structure in lipid environment. Structural comparison with the inactive conformation of H_2_R and the inactive and G_q_-coupled active state of H_1_R together with structure-guided functional experiments reveal molecular insights into the specificity of ligand binding and G protein coupling for this receptor family. We demonstrate lipid-modulated folding of cell-free synthesized H_2_R, its agonist-dependent internalization and its interaction with endogenously synthesized H_1_R and H_2_R in HEK293 cells by applying a recently developed nanotransfer technique.

## Introduction

The biogenic amine histamine is a hormone and neurotransmitter that is ubiquitously distributed in the human body. It plays central roles in diverse (patho)physiological processes, such as inflammation, allergy, gastric acid secretion, cellular migration, vasodilatation, bronchoconstriction and neurotransmission^1, 2^. Histamine signaling is mediated through binding and activation of the histamine 1-4 receptor subtypes H_1_R, H_2_R, H_3_R and H_4_R belonging to the family of G protein-coupled receptors (GPCRs)^1^. The H_2_R is an important regulator of gastric acid secretion, ionotropic and chronotropic cardiac stimulation, vasodilatation and mucus production^1, 3^. Histamine binding to the H_2_R increases both, cAMP and inositol phosphate second messengers, by signaling through the heterotrimeric G proteins G_s_ and G_q_, respectively^4–6^. Clinical drugs targeting H_2_R such as famotidine, nizatidine and cimetidine are applied to suppress gastric acid secretion in esophageal reflux disease and are important for peptic, gastric and duodenal ulcer healing^1, 5, 7^. More recently, H_2_R is discussed as therapeutic target for several cardiovascular conditions as well as for acute myeloid leukemia, diabetes and colorectal cancer^8, 9^. The proposed H_2_R homodimerization and agonist-dependent cross-desensitization, co-internalization and heterodimerization with the H_1_R may provide additional routes for new therapeutic strategies^10^. A central challenge, however, has been the development of subtype-selective ligands with low off-target side effects. Thus, a more detailed structural basis of ligand binding specificity and of H_2_R activation is required to design drugs that can precisely tune H_2_R signaling.

Cell-free (CF) expression enables the production of proteins in presence of stabilizing ligands and it allows the direct insertion of nascent membrane proteins into defined membrane environments. The CF synthesis of GPCRs was continuously improved during the last decade^11–13^. In particular, the strategy to insert nascent GPCRs cotranslationally into nanodiscs (NDs) avoids any contacts to potentially denaturing detergents^11, 14–16^. The insertion into empty ND membranes is translocon independent and is accompanied by a release of lipids^17^. The lipid composition of ND membranes can be of crucial importance for the efficiency of cotranslational membrane insertion as well as for the function of the inserted membrane proteins^13, 16, 18^. The additional presence of ligands as well as of interacting G-proteins can further stabilize CF synthesized GPCRs and support their functional folding. By using these unique technical advantages, the formation of stable GPCR/G-protein complexes in CF reactions was recently demonstrated for the first time^13^.

Here, we used CF expression of the human H_2_R with *E. coli* lysates to determine the cryo electron microscopy (cryo-EM) structure of the receptor in its active conformation and in complex with the G_s_ heterotrimer at a global resolution of 3.4 Å. The complex was formed with full-length H_2_R, G_s_ and the G_s_ stabilizing nanobody Nb35 in ND membranes composed of DOPG lipid. The complex represents the first structure of a CF synthesized GPCR as well as the first GPCR/G_s_ complex structure obtained in lipid environment. The biochemical characterization is complemented by analysis of the CF synthesized H_2_R after transfer into membranes of HEK293T cells using a recently developed nanotransfer technique^19–21^.

## Results

### CF expression optimization and histamine/H_2_R/G_s_/Nb35/ND complex preparation

Full-length human H_2_R was cotranslationally inserted into preformed NDs by CF expression. Lipid screens revealed NDs containing the negatively charged lipids DOPG or DEPG as most efficient for H_2_R insertion (Extended Data Fig. S1A). SEC analysis of purified H_2_R/ND complexes indicated best sample quality with ND membranes composed out of DOPG, DEPG and DMPG (Extended Data Fig. S1B). Effects of various supplied ligands on cotranslationally synthesized H_2_R/ND (DOPG) complexes were then monitored by SEC profiling (Extended Data Fig. S1C). Stabilizing effects of the proposed folded H_2_R/ND fraction were observed after synthesis in presence of the agonist histamine and with most antagonists (Extended Data Fig. S1C).

Cryo-EM samples of H_2_R in the active state and complexed to the heterotrimeric G protein G_s_ were then produced by CF synthesis of H_2_R in presence of the agonist histamine, heterotrimeric G_s_ protein previously purified from insect cells, preformed NDs (DOPG) and the Nb35 containing a C-terminal His-tag (Nb35-His) in a total reaction volume of 1.8 mL (Fig. 1). After incubation for approx. 16 h, the reaction was treated with apyrase and the synthesized H_2_R/G_s_/Nb35-His complexes in NDs (DOPG) were purified by IMAC and subsequently analyzed by SDS-PAGE (Extended Data Fig. S2A+B) and SEC analysis (Extended Data Fig. S2C). The peak SEC fraction was taken for cryo-EM analysis and concentrated to 2.8 mg/mL in a final volume of 35 µL.

**Fig. 1.**
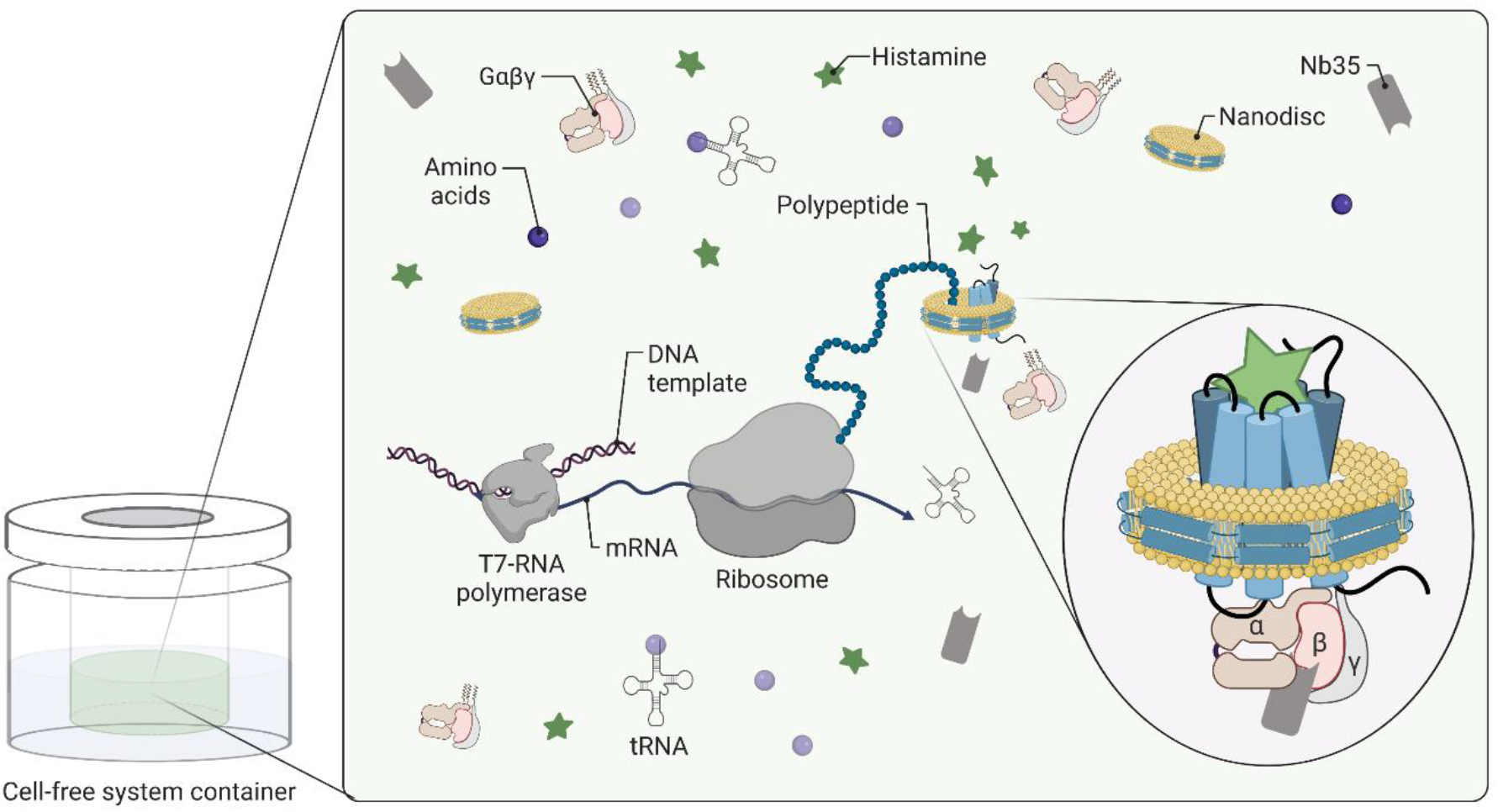
Cryo-EM sample preparation of complexes by CF expression. H_2_R is synthesized by cotranslational insertion into preformed NDs (DOPG) in presence of histamine and G_s_ heterotrimer purified from insect cells (created by biorender).

### Nanotransfer and oligomerization of CF synthesized H_2_R in HEK293T cells

The functionality of CF synthesized H_2_R was analyzed using a recently developed technique for GPCR transfer from ND membranes into membranes of living cells^19, 20^ (Fig. 2A). Ligand binding could be demonstrated by the histamine dependent internalization of transferred H_2_R-mNG derivatives in HEK293T cells via increase of the cytosolic fluorescence (Fig. 2B-D). Moreover, some co-localization of internalized transferred H_2_R-mNG with H_1_R-mCherry synthesized after transfection of the HEK293T cells indicate that receptors from the different origins take the same route of internalization.

**Fig. 2.**
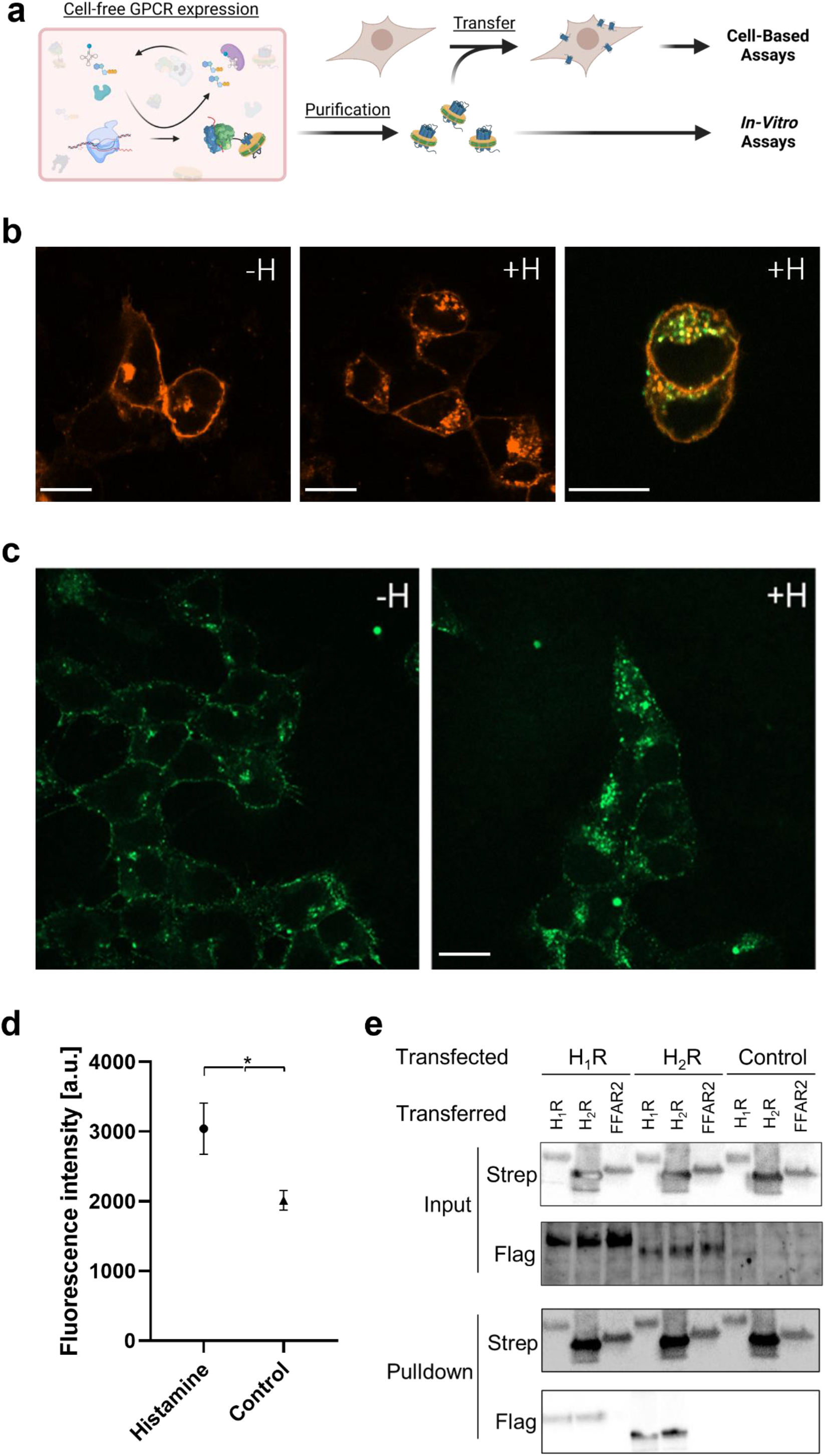
Characterization of CF synthesized H_2_R after nanotransfer into HEK293T cells. **a.** Schematic of GPCR nanotransfer. **b.** H_1_R-mCherry synthesized after transfection internalizes upon histamine treatment (100 µM histamine for 1h; + H) and co-localizes with nanotransferred and CF synthesized H_2_R-mNG (right panel). White bar = 10 µm. **c.** Representative images of HEK293T cells transferred for 4 h with 0.5 µM purified H_2_R-mNG in NDs (DOPG) and treated with histamine. Left panel: Localization of transferred H_2_R-mNG in untreated cells. Right panel: Internalization of transferred H_2_R after incubation with 100 µM histamine for 1h. White bar = 10 µm. **d.** Quantification of cytosolic fluorescence of transferred H_2_R-mNG. Data are presented as mean s.d. (n = 3 individual experiments, *p < 0.05, students t-test). **e.** Interaction of transferred CF synthesized GPCRs with transfected H_1_R and H_2_R. Transfected cells expressing Flag-tagged H_1_R or H_2_R were transferred with CF synthesized Strep-tagged H_1_R, H_2_R or FFAR_2_ for 4 h. Subsequently, cells were washed, lysed and transferred GPCRs were immobilized by anti-Strep pulldowns. Input: Immunoblots of cell lysates before anti-Strep pulldowns. Pulldown: Immunoblots of immobilized GPCRs after anti-Strep pulldowns. Strep: Anti-Strep antibody; Flag: Anti-Flag antibody; Control: Cells without transfection (a. created by biorender).

H_2_R is known to form stable homo-oligomers, in addition to hetero-oligomerization with the related receptor H_1_R^10, 22, 23^. To study these interactions, CF synthesized GPCRs containing a C-terminal Strep-tag were transferred into HEK293T cells that were previously transfected with expression vectors encoding for Flag-tagged GPCR derivatives (Fig. 2E). Strep-tagged H_2_R, Strep-tagged H_1_R or Strep-tagged free fatty acid 2 receptor (FFAR_2_) serving as negative control were cotranslationally inserted into NDs (DOPG), purified and transferred into HEK293T cells previously transfected with constructs encoding for Flag-H_1_R or Flag-H_2_R. After washing and cell lysis, the transferred GPCRs were immobilized by anti-Strep antibodies and co-immobilized GPCRs were identified by anti-Flag antibodies (Fig.2E). The results revealed both, homodimerization of H_1_R and H_2_R as well as the H_1_R-H_2_R heterodimerization, while no interactions of transferred Strep-tagged FFAR_2_ with any of the histamine receptors was detected (Fig. 2E). These data agree with previous reports on the oligomerization of the tested GPCRs and further indicate the correct folding of the CF synthesized GPCRs after their transfer in HEK293T cells.

### Structure determination of the Histamine/H_2_R/G_s_ signaling complex

We determined a cryo-EM structure of the histamine-bound H_2_R/G_s_ complex stabilized by Nb35 and embedded in ND (DOPG) membranes at a global resolution of 3.4 Å (Fig. 3 and Extended Data Fig. S3). Representative class averages and data analysis flow chart are shown in Extended Data Fig. S4, associated statistics is summarized in Extended Data Table S1. The cryo-EM maps show an evenly distributed resolution with strong density for the bound agonist histamine and for the TM domains of H_2_R (Extended Data Fig. S5). The cryo-EM map enabled building of an atomic model for the histamine-bound active conformation of H_2_R (residues D13^1.27^-G298^8.53^) (superscript numbers indicate generic GPCR numbering following the revised Ballesteros-Weinstein system for family A GPCRs) including the canonical seven transmembrane helices (TMs 1-7) connected by three extracellular loops (ECLs 1-3) and three intracellular loops (ICL1-3), a common architecture of GPCRs (Extended Data Fig. S6). Due to the conformational heterogeneity of the C-terminal end of helix 8 (H8) and the resulting weak density, the receptor was only modeled until residue G298^8.53^. The increased structural flexibility in this region could be a consequence of the missing palmitoylation of residue C304^8.59^ at the C-terminal end of H8 of the CF-expressed receptor that is predicted to anchor H8 to the lipid membrane^24–26^. Otherwise, the local resolution map shows a relatively even distributed quality across the entire receptor complex with weaker density for the central part of ECL2, which remains partially unresolved (residues T164^ECL^^2^-T171^ECL^^2^) most likely due to its conformational flexibility. The C-terminal region of ECL2, however, is stabilized by one conserved disulfide bond connecting residue C91^3.25^ in the N-terminal end of TM3 with residue C174^45.40^ in ECL2.

**Fig. 3.**
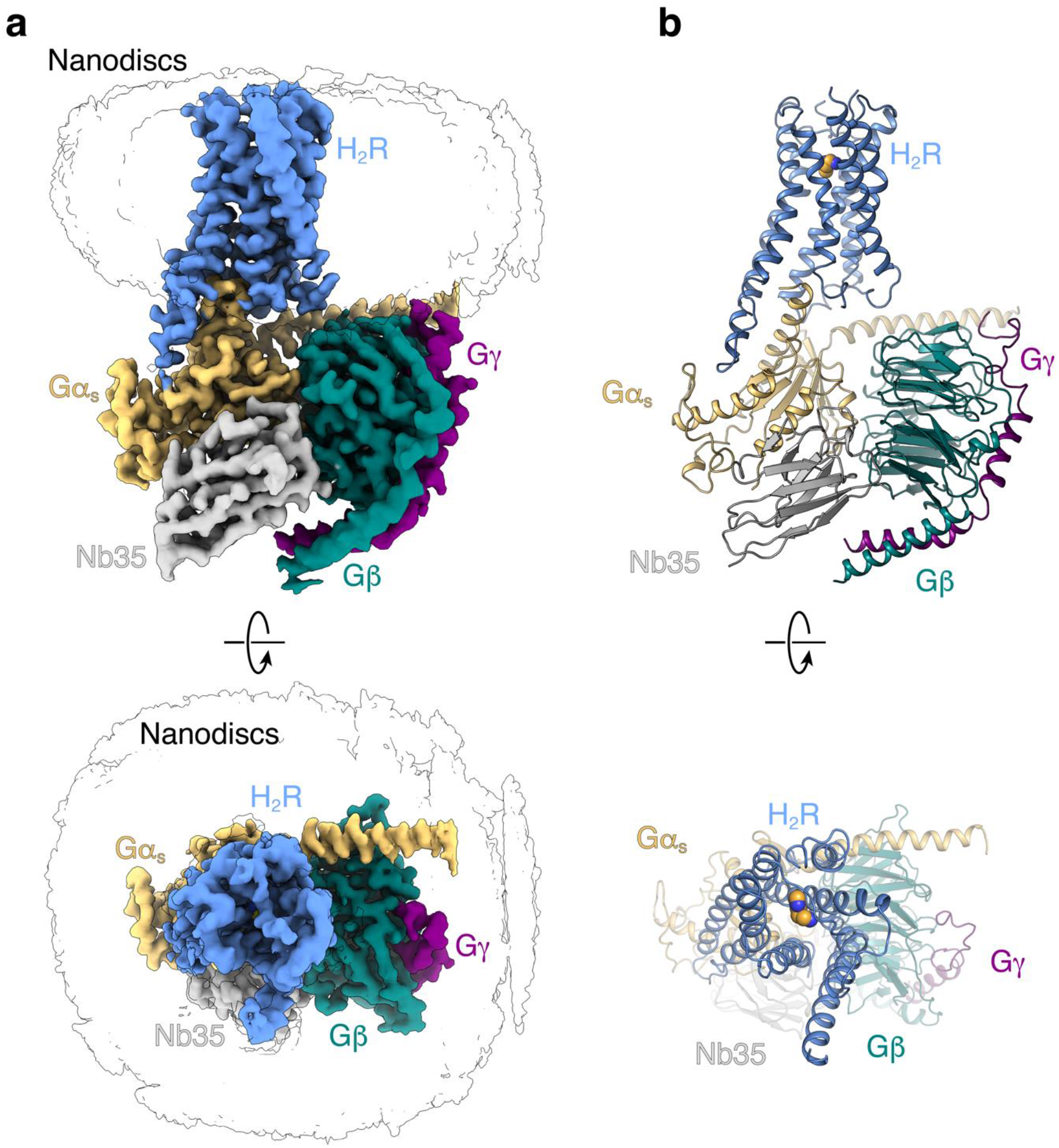
Cryo-EM structure of the histamine-bound H_2_R-Gs/Nb35/ND complex. **a.** Cryo-EM density map of the histamine-bound H_2_R-Gs/Nb35/ND complex in nanodiscs colored by subunit. Blue, H2R; wheat, Gαs; cyan, Gβ; purple, Gγ, grey, Nb35; black silhouettes, nanodiscs. **b.** Ribbon model of the H_2_R-Gs/Nb35/ND complex bound to histamine (yellow spheres) colored by subunit.

### Comparison of histamine binding between the H_2_R and H_1_R

The bound histamine shows clear density within the orthosteric ligand binding pocket of H_2_R formed by transmembrane domains (TMs) 3, 5, 6, and 7 (Fig. 4, Extended data Fig. S5). The primary amine of histamine (N^α^) forms H-bonds and ionic interactions with the backbone carbonyl and the carboxyl group of D98^3.32^ in TM3, respectively — a highly conserved residue among aminergic receptors. Another H-bond interaction is formed between the N^π^ atom of the imidazole ring and the conserved Y250^6.51^ in TM6 (Fig. 4A, Extended data Fig. S6). The rest of the histamine binding pocket is composed of hydrophobic aliphatic or aromatic residues in TM3 (V99^3.33^) and TM6 (Y250^6.51^, F251^6.52^, F254^6.55^, and W247^6.48^) as well as polar side chains in TM3 (T103^3.37^), TM5 (D186^5.42^, T190^5.46^) and TM7 (Y278^7.43^).

**Fig. 4.**
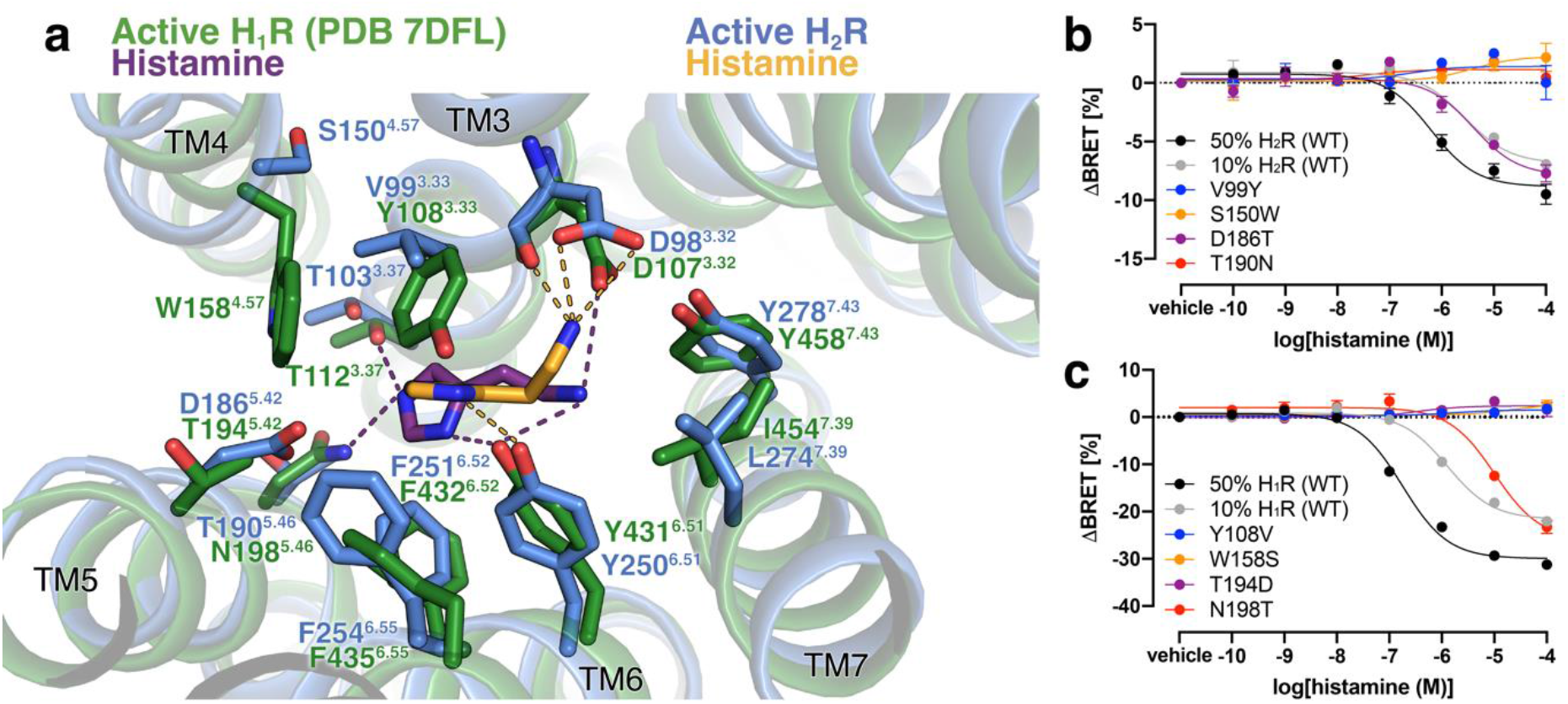
Histamine-binding to H_2_R. **a.** Comparison of the histamine-binding site in H_2_R (blue, receptor; yellow, histamine) and H_1_R (green, receptor; purple, histamine) (PDB ID 7DFL). Binding pocket residues are represented as sticks and labeled with residue number and Ballesteros-Weinstein code (superscript) colored by the receptor subtype. H-bond interactions are shown as dotted lines colored by the interacting ligand. **b.** and **c.** Mutagenesis analysis of receptor-specific residues within the histamine-binding pocket in H_2_R (**b**) and H_1_R (**c**) using a BRET-based G protein dissociation assay. Signaling graphs show mean ± s.e.m. of three independent biological replicates with a global fit of the data.

Comparison of the H_2_R-G_s_ structure with the available active structure of the H_1_R-G_q_ complex (PDB ID 7DFL)^27^ revealed some receptor subtype specific characteristics (Fig. 4A). As a common feature, the nitrogen atoms N^α^ and N^π^ of the bound histamine engage in polar interactions with residues D^3.32^ and Y^6.51^, respectively. However, amino acid sequence variations in the orthosteric ligand binding site between the two receptor subtypes cause a rotation of the histamine imidazole ring in the H_2_R by approximately 80° with respect to the ligand binding pose in the H_1_R. The rotation prevents a clash of the side chain of Y^3.33^ in H_1_R (V^3.33^ in H_2_R) with the perpendicular oriented imidazole ring in the H_2_R structure and causes an overall deeper binding position towards the H_1_R receptor core compared to the H_2_R. As a result, the N^t^ atom of histamine is able to form H-bonds with T112^3.37^ in TM3 and N198^5.46^ in TM5 of H_1_R. In contrast, the distances between the N^t^ atom of the histamine and the corresponding residues T103^3.37^ in TM3 and T190^5.46^ in TM5 of H_2_R are too large (4.9 Å and 6.5 Å, respectively) to allow hydrogen bonding interactions. Furthermore, the non-conserved residue D186^5.42^ in TM5 of the H_2_R is not in H-bonding distance to the ligand and therefore does not directly contribute to histamine-receptor interactions. This agrees with previous functional studies showing that this residue is crucial for H_2_R-specific antagonist, but not histamine binding^28^.

Based on these observations, subtype-specific residues in the H_2_R and H_1_R were tested by mutagenesis for their contribution to histamine-dependent receptor-mediated G protein activation. Residues at positions 3.33, 4.57, 5.42 and 5.46 in H_2_R were individually mutated to the corresponding residues in H_1_R and *vice versa,* and functionally characterized using bioluminescence resonance energy transfer (BRET)-based G protein activation (G-CASE) sensors^29^ (Fig.4B). All H_1_R mutations maintained >40% of wild-type (WT) surface expression levels, while the expression levels of the H_2_R mutations S150W^4.57^, D186T^5.42^, and T190N^5.46^ were significantly reduced or almost completely abolished (Extended Data Fig S7). Therefore, transfected DNA of the WT H_2_R was adjusted accordingly to allow comparison at similar expression levels. The H_2_R mutations V99Y^3.33^, S150W^4.57^, and T190N^5.46^ completely ablated H_2_R-mediated G_s_ activation. Consistent with the H_2_R-G_s_ complex structure and the functional studies mentioned above^28^, D186T^5.42^ had no impact on histamine-dependent receptor signaling. While low receptor amount in the plasma membrane might contribute to the missing activity of the T190N^5.46^ and S150W^4.57^ mutants, the effect of the V99Y^3.33^ and S150W^4.57^ mutations are presumably due to the larger size of the introduced tyrosine and tryptophan side chains. In particular, the substitution of V99^3.33^ by tyrosine might result in steric hindrance of histamine binding in the H_2_R, while a bulky tryptophan side chain at position 4.57 in TM4 presumably causes distortion of the ligand binding pocket due to clashes with residues in TM3. In the H_1_R, mutations Y108V^3.33^, W158S^4.57^, and T194D^5.42^ completely ablated receptor-mediated G_q_ activation, whereas the substitution N198T^5.46^ led to an 8-fold lower histamine potency (Fig. 4C). Besides the conserved common polar contacts between histamine and residues D^3.32^ and Y^6.51^ in all histamine receptors, these results confirm that the H_1_R and H_2_R exhibit subtype-specific receptor-ligand interactions that may also contribute to the molecular basis of agonist-binding selectivity.

### H_2_R activation by histamine

Compared to the inactive structure of the H_2_R bound to the antagonist famotidine (PDB ID 7IL3)^30^, the orthosteric ligand binding pocket in histamine-bound H_2_R undergoes some subtle structural rearrangements caused by differences in receptor-ligand interactions (Fig. 5). Specifically, the H-bond between the imidazole nitrogen N^π^ of the agonist and Y^6.51^ in TM6, which is absent in the famotidine-bound inactive structure, seems to cause an inward movement of the extracellular end of TM6 (1.8 Å as measured between the C_α_ atoms of G258). Furthermore, the 3.8 Å downwards shift of the primary amine group of the agonist towards the intracellular side in comparison to the corresponding amine of the antagonist enables the formation of stronger H-bond interactions of the agonist with the backbone and side chain of the conserved D98^3.32^ in TM3. This interaction might result in the observed counter-clockwise rotation of TM3 upon receptor activation. Another activation-dependent conformational change on the extracellular side of the receptor involves the slight movement of TM5 towards TM4. Notably, while histamine does not engage TM5, the guanidinium group of famotidine forms H-bond interactions with residues D186^5.42^ and T190^5.46^ in TM5 as well as T103^3.37^ in TM3, which presumably prevents movement of these transmembrane helices and stabilizes the receptor in the inactive state.

**Fig. 5.**
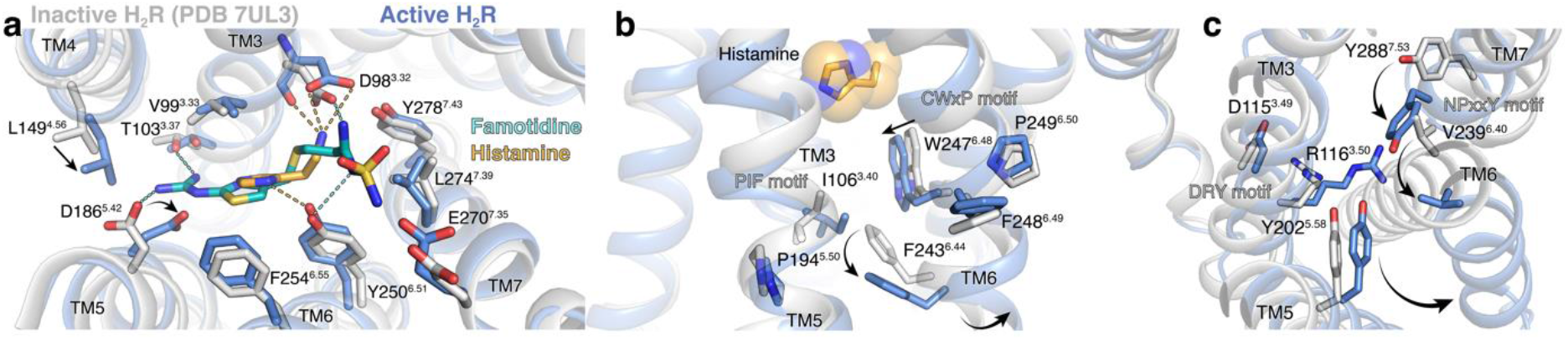
H_2_R activation by histamine. **a.** Binding pocket of the histamine-activated H_2_R (blue) with the inactive famotidine-bound receptor (gray) (PDB ID 7UL3) overlaid. H-bond interactions are shown as dotted lines colored by the interacting ligand. **b.** Close-up view of the CWxP and PIF motif in the inactive (grey) H_2_R and the active (blue) H_2_R bound to histamine (yellow spheres). **c.** Comparison of the DRY motif and the NPxxY motif in the inactive (grey) H_2_R and active (blue) H_2_R structures. Amino acid side chains are represented as sticks and labeled with residue number and Ballesteros-Weinstein code (superscript) colored by the receptor subtype. Arrows indicate conformational changes in the H_2_R upon activation.

More pronounced conformational changes take place on the intracellular side typical for the activation of family A GPCRs (Fig. 5)^31^. This includes the outward rotation of the cytoplasmic end of TM6 and concerted movements of the intracellular end of TM5 towards TM6 as well as the inward shift of the intracellular part of the adjacent TM7, resulting in the creation of the intracellular G protein-binding site (Fig. 5). Previous studies have identified conserved sequence motifs in family A GPCRs, also known as microswitches that are important for transmitting the ligand-induced conformational changes from the orthosteric ligand binding pocket to the G protein-binding cavity^31, 32^. Comparison of structural alterations in those microswitch regions between the inactive and active structures of the H_2_R, the H_1_R and the β_2_ adrenergic receptor (β_2_AR) revealed nearly identical rearrangements, suggesting that these aminergic receptors share a similar activation mechanism (Extended Data Fig. S8). In particular, the ‘toggle-switch’ residue W^6.48^ in the C^6.47^W^6.48^xP^6.50^ motif at the bottom of the orthosteric ligand binding pocket is displaced upon receptor activation. Nearer to the intracellular side of the receptor, the P^5.50^I^3.40^F^6.44^ motif undergoes a ratchet like motion that includes an outward movement of F^6.44^ past residue I^3.40^. Altogether, these conformational changes eventually induce the outward rotation of TM6 to enable opening of the intracellular G protein-coupling cavity and subsequent engagement of the heterotrimeric G protein. The formation of the active state is further facilitated by rearrangements of the highly conserved D^3.49^R^3.50^Y^3.51^ motif and the N^7.49^P^7.50^xxY^7.53^ motif on the intracellular side of TM3 and TM7, respectively.

### Agonist binding selectivity of H_2_R and H_1_R

H_1_R and H_2_R appear to have a conserved histamine recognition motif involving main contacts to residues D^3.32^ and Y^5.51^, whereas variation in the interaction with TM5 at position 5.42 or 5.46 may have a major contribution to agonist selectivity. To investigate the molecular basis of selective binding of H_2_R-specific agonists, we first analyzed predicted binding modes by generating a variety of possible histamine orientations in the H_2_R using SEED (Extended Data Fig. S9A). Considering the predicted binding energy of each pose, the canonical histamine orientation was indeed the top-scored one. Comparison of further predicted histamine binding modes in both receptors with the respective experimental ones showed that the docking software AutoDock Vina generated best matching poses with a root-mean-square deviation (r.m.s.d.) of the heavy atom positions of 0.76 Å for the H_1_R and 0.57 Å for the H_2_R (Extended Data Fig. S9B).

AutoDock Vina was then used to predict the binding modes for various H_2_R-selective agonists in both receptors. The agonist amthamine engages in charge-assisted H-bond interactions with the conserved residue D98^3.32^ as well as with the H_2_R-specific residue D186^5.42^ (Fig. 6 A+B). In contrast, in the H_1_R, amthamine is predicted in a different orientation and interacts with D107^3.32^ via the uncharged amino group, an interaction that lacks the ionic character, and has no direct contact with T194^5.42^ in TM5 (D186^5.42^ in the H_2_R). In addition, the methyl group of amthamine is predicted to engage in a favorable hydrophobic interaction with the side chain of F251^6.52^ in the H_2_R that is not observed in the H_1_R. Moreover, the presence of the bulky residue Y108^3.33^ in TM3 of the H_1_R seems to prevent any of the investigated H_2_R-selective agonists from adopting a binding mode that is similar to the respective ones found in the H_2_R (Fig. 6 B).

**Fig. 6.**
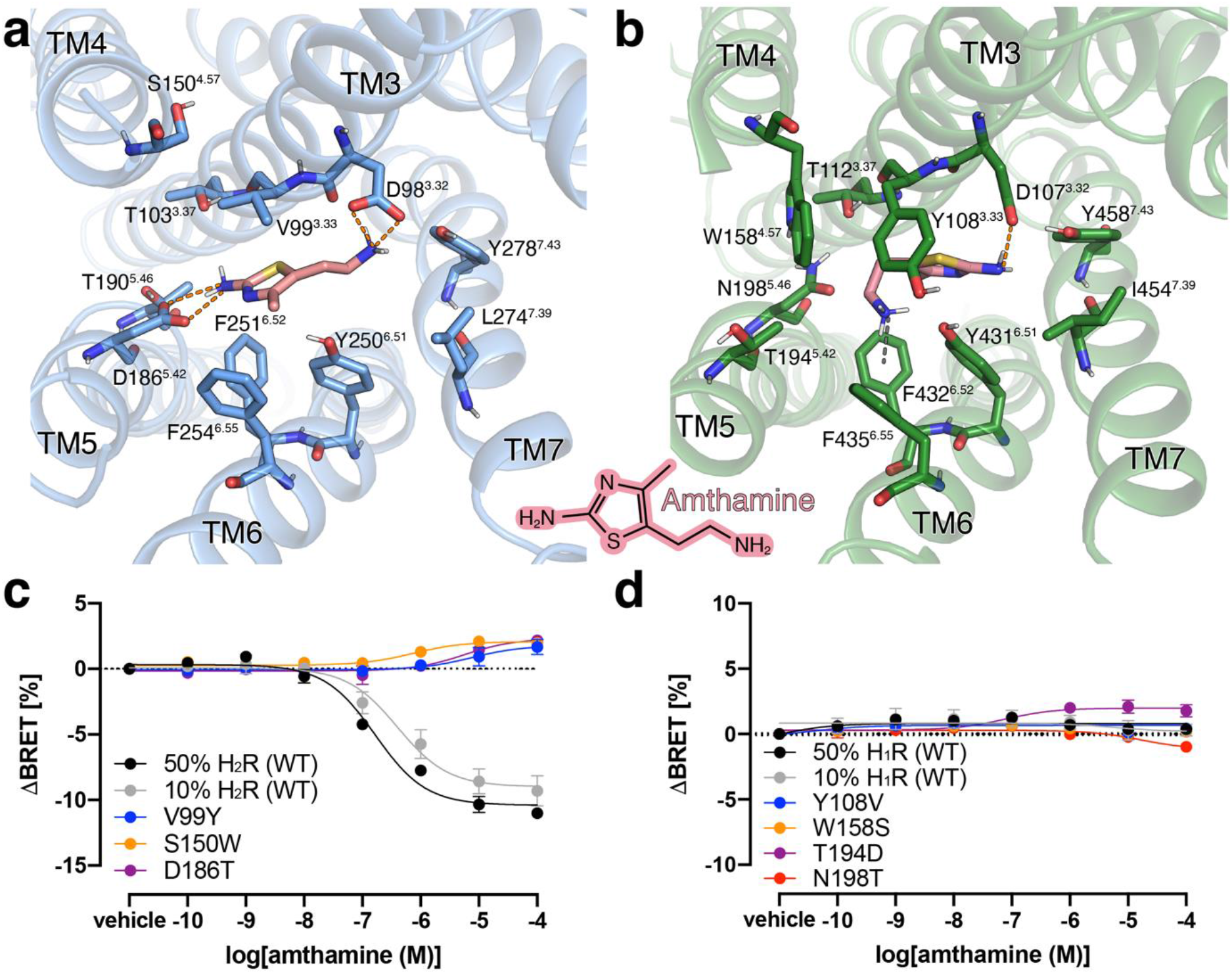
Agonist binding selectivity of H_2_R and H_1_R. **a.** and **b.** Predicted binding modes for the H_2_R selective agonist amthamine (light pink) in (**a**) H_2_R (blue) and (**b**) H_1_R (green). The orange dotted lines represent H-bond interactions. The green dotted line represents a possible cation-aromatic interaction between F432^6.52^ in H_1_R and the protonated amino group of amthamine. Binding pocket residues are represented as sticks and labeled with residue number and Ballesteros-Weinstein code (superscript). Binding mode predictions were obtained with AutoDock Vina. **c.** and **d.** Mutagenesis analysis of receptor-specific residues within the ligand-binding pocket in (**c**) H_2_R and (**d**) H_1_R using a BRET-based G protein dissociation assay in the presence of amthamine. Signaling graphs show mean ± s.e.m. of three independent biological replicates with a global fit of the data.

To experimentally validate the predicted complexes, we used H_1_R and H_2_R mutants that were designed by swapping the receptor-specific amino acids in TM3-TM5, as described above, and performed BRET-based signaling assays in comparison to the WT receptors in the presence of amthamine (Fig. 6 C+D). Notably, the H_2_R mutant T190N^5.46^ was not included in these experiments because of its very low expression level and activity (Extended Data Fig. S7). Similar to histamine (Fig. 4 B+C), the H_2_R mutations V99Y^3.33^ and S150W^4.57^ completely abolished amthamine-stimulated G_s_ activation and supported the proposed steric hindrance of agonist binding. However, signaling of the mutant H_2_R-D186T^5.42^ was remarkably different to that of histamine as it was completely unresponsive to amthamine (Fig. 6C). This inactivation is most likely caused by disrupting the interaction between D186^5.42^ and the aromatic amine on the amthamine thiazole ring. For the H_1_R WT and mutants, we observed no or only marginally amthamine-dependent stimulation of G_q_ signaling (Fig. 6D).

In addition to amthamine, possible binding modes in the H_1_R and H_2_R were also predicted for the H_2_R-selective agonists dimaprit and the carbamoylguanidine-derivative, compound 157^33^ (reported K_i_ selectivity H_2_R/H_1_R ratio of 1:3802) (Fig. 7 and Extended Data Tables S2 and S3). As in the case with amthamine, the interaction with D186^5.42^ can be identified as the main predicted reason for H_2_R selectivity of both compounds. In summary, these functional studies in combination with the predicted binding modes of H_2_R selective agonists underline the importance of positions 5.42 and 5.46 not only for selective binding of H_2_R blockers as shown previously^28^, but also for the investigated H_2_R agonists.

**Fig. 7.**
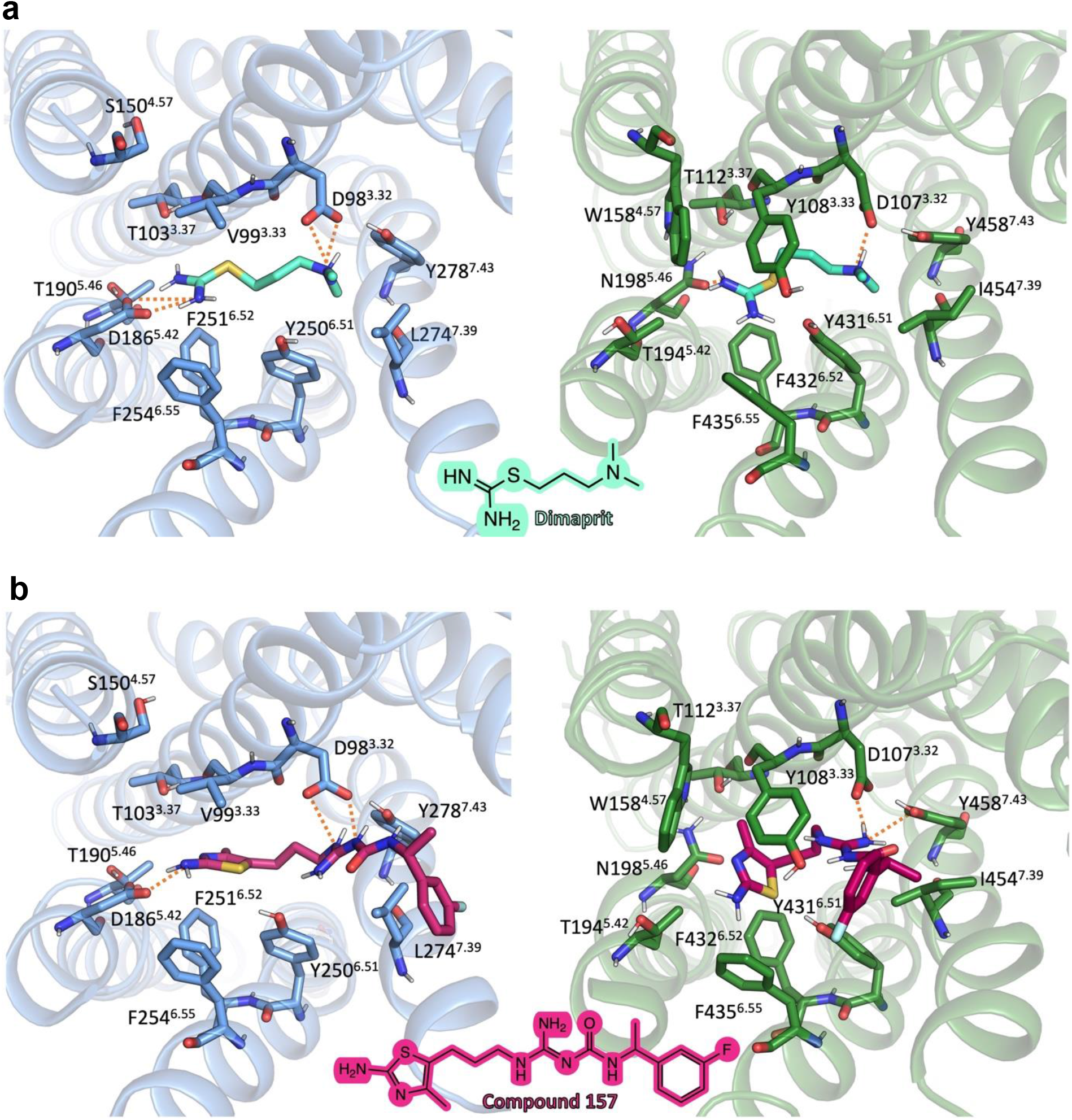
Molecular docking of H_2_R specific agonists in the H_2_R and the H_1_R. **a.** Docking of dimaprit; **b.** Docking of compound 157^33^. Structural elements and relevant residues of the binding pockets of the H_2_R (blue) or the H_1_R (green) are shown. The orange dotted lines represent H-bond interactions. Binding pocket residues are represented as sticks and labeled with residue number and Ballesteros-Weinstein number (superscript). Binding mode predictions were obtained with AutoDock Vina.

### G protein coupling to the H_2_R

The overall structure of the histamine-activated H_2_R-G_s_ complex in a lipid ND shows a similar receptor-G protein conformation compared to other available aminergic receptor-G_s_ complex structures obtained in detergent (r.m.s.d. of 0.9 to 2.7 Å) (Extended Data Fig. S10.). While all aminergic receptor-G_s_ complexes display the prototypical receptor-G protein conformation, they also exhibit significant differences in the relative orientation of the coupled G protein with respect to the receptor. Specifically, the N-terminal αN helix of the Gα_s_ subunit is rotated up to approximately 35° in the membrane plane relative to the TM bundle between different receptor-G protein complexes. In the H_2_R-G_s_ complex, the relative orientation of the αN helix is most similar to the one found in the complex structures of the adrenergic receptor β_2_AR (PDB IDs 3SN6, 7BZ2, 7DHI, 7DHR), and the dopamine receptor D_1_R (PDB ID 7F1Z) (Extended Data Fig. S10B). The position of the αN helix of the H_2_R-coupled Gα_s_ subunit with respect to the membrane seems to be stabilized by three lysine (K8, K17, and K24) and two arginine residues (R13 and R20) that are oriented towards the membrane and may interact with the negatively charged polar headgroups of the POPG lipids in the ND (Extended Data Fig. S10C). Notably, similar electrostatic interactions between basic residues in the αN helix and the lipid bilayer have been found in G_i_ complex structures of the dopamine receptor and the neurotensin receptor 1 NTSR1^34^ in NDs, demonstrating that these N-terminal polybasic regions play an important role for membrane anchoring of different G protein families in agreement with previous mutagenesis and functional studies^35, 36^.

The interface between the H_2_R and G_s_ includes a buried surface area of 2951 Å^2^. Like in other aminergic receptor-G_s_ complexes, the main interaction sites occur between ICL2 and TM3, TM5, TM6 and the TM7-H8 kink on the receptor and the αN-β1 hinge loop and α5 on the Gα subunit of the G protein. Upon coupling to the H_2_R, the α5 helix of the G protein rotates by 60° and straightens up due to a 13 Å translation of its distal C-terminal end towards TM6 in comparison to the GDP-bound inactive structure of G ^37^. This movement is accompanied by a 5 Å shift of α5 towards the receptor to form a number of hydrophobic and hydrophilic contacts with residues in the intracellular G protein binding cavity of the H_2_R. Together with the interaction between ICL2 of the receptor and the αN-β1 hinge loop and α5 helix of the G protein, which has been shown to impact the conformational dynamics of the β1 strand and the adjacent nucleotide-binding P loop of the Gα_s_ subunit in complex with the β_2_AR receptor^38^, the rotational translation of the α5 helix leads to a disruption of the GDP-binding pocket and subsequent nucleotide release.

### Differences between the H_2_R-G_s_ and H_1_R-G_q_ G protein-coupling interface

When the structures of the H_2_R-G_s_ and the H_1_R-G_q_ complexes are superimposed on the receptor, the G_s_ and G_q_ proteins display differences in their orientation relative to the TM bundle (Fig.8A). The Ras domain of Gα_s_ is shifted and slightly rotated away from TM3 towards TM6 when compared to the one of Gα_q_. This difference is propagated to the αN helix and the Gβγ subunits of the G protein, resulting in significant differences in the receptor-G protein interactions relative to the H_1_R-G_q_ complex. In particular, the α5 helix is more shifted towards the TM5/TM6 region of the H_2_R than in the H_1_R-G_q_ complex. This closer position of α5 of Gα_s_ relative to TM5 and TM6 presumably leads to the slightly further outward displacement of TM6 in the H_2_R compared to the H_1_R (Fig. 8A). Furthermore, the tighter interaction between α5 and TMs 5 and 6 allows the formation of a H-bond interaction between the backbone carbonyl of E392 of Gα_s_ and the side chain of T235^6.36^ as well as potential polar interactions of D381, Q384, and R385 of the G protein with residues in TM5 (Q212^5.68^ and R215^5.71^) (Fig. 8B). Together with the van der Waals contacts formed between residues Y358 and I216^5.72^ as well as L346 and I219^ICL3^ of Gα_s_ and the receptor, respectively, these interactions putatively stabilize the more extended α-helical structure of the C-terminal end of TM5 in comparison to the TM5 of H_1_R, which forms weaker contacts in this region with the engaged Gα_q_ (Fig. 8C).

**Fig. 8.**
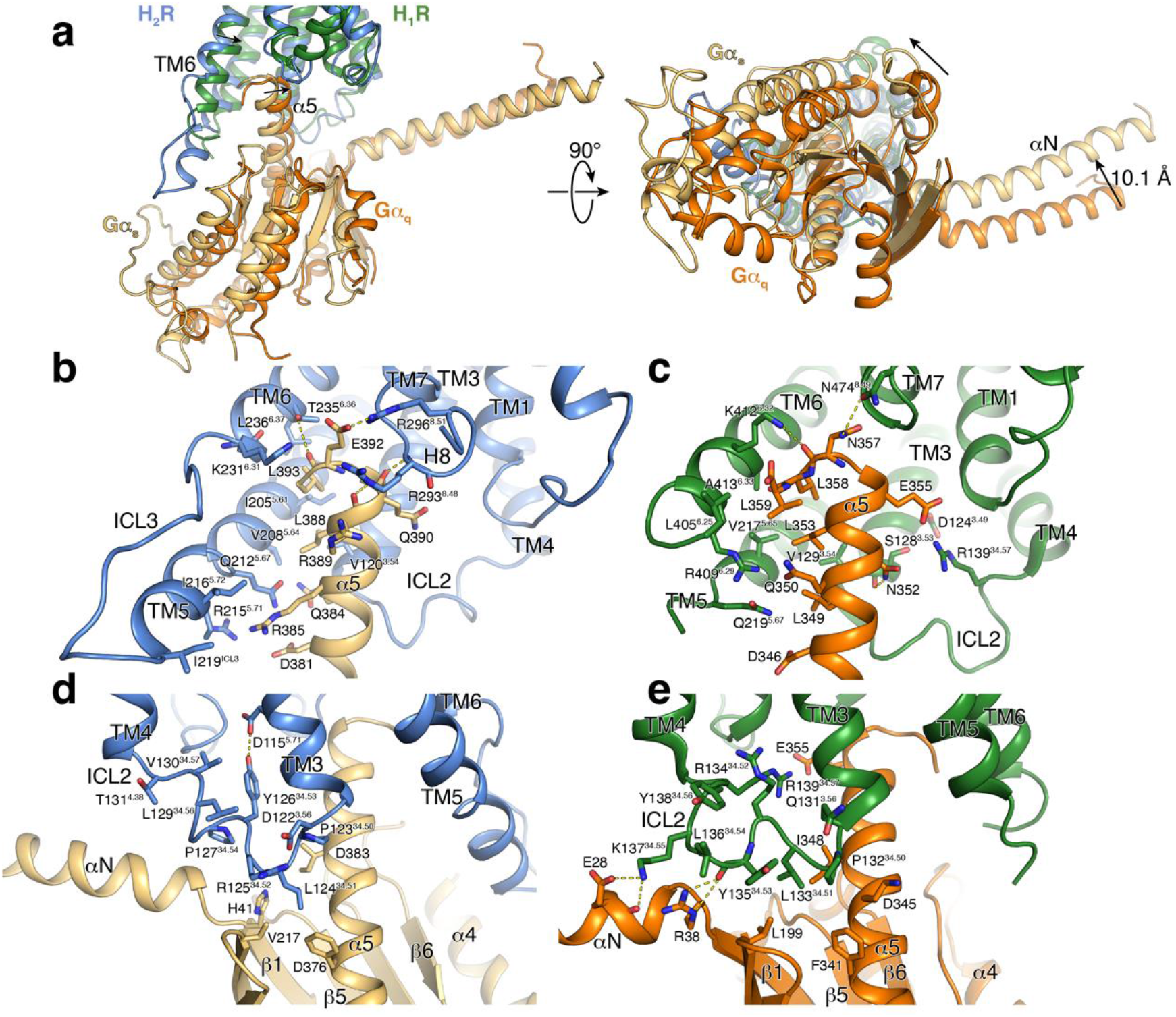
Comparison of G protein interfaces between the H_2_R-G_s_ and the H_1_R-G_q_ complex. **a.** Comparison of the relative receptor-G protein orientation between the H_2_R (blue) and the H_1_R (green) subtype coupled to Gα_s_ (wheat) and Gα_q_ (orange), respectively. **b.** and **c.** Comparison of the interface between (**b**) the α5 of Gα_s_ and H_2_R and (**c**) the α5 of Gα_q_ and H_1_R. **d.** and **e.** Comparison of the interface between the ICL2 and the G protein subtype in (**d**) the H_1_R-Gα_s_ complex and (**e**) the H_1_R-Gα_q_ complex. Interface residues are represented as sticks and labeled with residue number and for the receptor residues with the Ballesteros-Weinstein code (superscript). Arrows indicate differences in the relative orientation of the G protein subtypes with respect to the receptors.

In addition to the H-bond between E392 of Gα_s_ and T235^6.36^ of the H_2_R already mentioned above, the C-terminus of the G protein α5 forms additional interactions with the TM5/TM6 region as well as the TM7/H8 loop and H8 of the H_2_R (Fig. 8B). While the very C-terminal residue L394 in the “hook” of the Gα_s_ α5 helix could not be modeled presumably due to its high flexibility, the adjacent conserved L393 is buried in a hydrophobic pocket lined by residues I205^5.61^, T235^6.36^, L236^6.37^ in TMs 5 and 6 of the H_2_R and L368 of the α5 helix. Furthermore, residues R389, Q390, and E392 of the G protein are engaged in H-bond interactions with two arginine residues, R293^8.48^ and R296^8.51^, located in the TM7/H8 hinge region and in H8 of the H_2_R, respectively. This interaction potentially causes the shift of the backbone of R293^8.48^ away from the receptor and closer to α5 in comparison to the H_1_R-Gα_q_ complex to engage in H-bonding with the Gα_s_. For Gα_q_, similar interactions are being formed between L358 and hydrophobic residues in TMs 5 and 6 of the H_1_R as well as N357 and N474^7.31^ in the TM7/H8 hinge region of the receptor. Overall, however, the α5 of Gα_q_ seems to engage in fewer interactions with TM5 and the TM7/H8 region of H_1_R compared to G_s_ with H_2_R (Fig. 7C).

Additional differences between the two complex structures are also observed in the ICL2-G protein interactions. Specifically, in ICL2 of the H_2_R-G_s_ structure, the conserved L124^34.51^ is positioned deeply inside the hydrophobic pocket formed by residues F376, I383 in α5 as well as V203 in β3 and H41 in β1 of Gα_s_ (Fig. 8D). As seen in other aminergic receptor-G_s_ complex structures, the ICL2 of H_2_R forms a helix that is stabilized by an interaction between the conserved residues Y126^34.53^ in the middle of the loop and D115^3.49^ of the conserved DRY motif in TM3. In contrast, ICL2 of the H_1_R adopts a position farther away from the Gα subunit so that L133^35.51^ does not reach as deep into the hydrophobic pocket between α5, β1 and β3 of Gα_q_ as in the H_2_R-G_s_ complex (Fig. 8E). Instead, the ICL2 of H_1_R forms H-bonds between K137^34.55^ and E28 and R134^34.52^ and D124^3.49^ of the DRY motif. The homologous residue to Y126^34.53^ in the H_2_R (Y135 in the H_1_R) does not interact with the DRY motif in the H_1_R-Gα_q_ structures but forms a H-bond interaction with its backbone carbonyl and R38 in the αN-β1 loop region of the G protein. In summary, these differences in the receptor-G protein interface might explain some of the underlying molecular principles of the G protein-coupling specificity observed for the H_1_R and the H_2_R histamine receptor subtypes^39–44^.

## Discussion

Here, we present the first report of a CF production pipeline to obtain a cryo-EM structure of a GPCR/G-protein complex. By using a semi-defined CF environment obtained from *E. coli* strain A19 in combination with supplied preformed lipid NDs, heterotrimeric G_s_ protein and Nb35, we were able to obtain a structure of the full-length H_2_R-G_s_ complex with its endogenous agonist histamine. The determined nominal resolution of 3.4 Å is comparable to analogous structures of GPCR complexes expressed and purified from eukaryotic cell-based systems^34, 45, 46^. In contrast to the classical cell-based approaches that require the detergent extraction of the GPCRs out of cell membranes, the presented CF approach uses the detergent-free cotranslational insertion of receptors directly into preformed ND membranes. The CF approach thus represents a shorter, milder and less complex strategy compared to cell-based procedures, avoiding any detergent solubilization and reconstitution procedures of GPCRs that may result in receptor denaturation^47, 48^. Furthermore, due to the simplified and translocon independent membrane insertion process, extensive engineering such as the deletion of terminal or internal domains is not necessary and full-length GPCR can be synthesized^12, 13, 49^.

Our combined strategy implementing cryo-EM, mutagenesis, signaling assays and docking studies yields important insights into the molecular underpinnings of ligand binding, receptor activation and G protein coupling of the H_2_R. Previous modelling studies using the β_2_-adrenoceptor/G_s_ complex^50^ as template and histamine insertion by molecular dynamics simulations^51^ proposed a salt bridge between the histamine ammonium group and residue D98^3.32^ of H_2_R, in addition to polar interactions between one imidazole nitrogen and residues D186^5.42^ and T190^5.46^. Our structure supports the interaction of the histamine ammonium group with D98^3.32^. However, distances of the histamine imidazole ring to residues D186^5.42^ and in particular to T190^5.46^ are too far for polar interactions. Instead, the binding of histamine is stabilized by an H-bond interaction between the ring’s N^π^ atom and Y250^6.51^. Besides these conserved key interactions between histamine and the conserved residues D^3.32^ and Y^6.51^ of histamine receptors, the divergent and less conserved residues at positions 5.42 and 5.46 in TM5 between the H_2_R and H_1_R were identified to participate in the ligand binding selectivity of the H_2_R. In the H_1_R, the histamine engages in H-bond interactions with N^5.46^, whereas in the H_2_R, the agonist does not form direct contacts with the corresponding residue T^5.46^ as well as the nearby H_2_R-specific residue D^5.42^. However, the latter residue seems to be important for the selective binding of H_2_R-specific agonists, as shown by our docking studies and by site-specific mutation of D5.42 to threonine, which specifically diminishes amthamine but not histamine signaling.

Furthermore, new insights into the molecular mechanism of H_2_R activation and on the differences in the receptor-G protein interactions between G_s_ and G_q_-coupled histamine receptor subtypes could be obtained. While the conserved microswitch motifs of the H_2_R undergo similar conformational changes as in the H_1_R, major differences are observed in the receptor-G protein coupling between the two receptors. The heterotrimeric G_s_ protein in the H_2_R-G_s_ complex is shifted more towards the TM 5/6 region of the receptor in comparison to G_q_ in the H_1_R-G_q_ complex. This translation results in the formation of tighter contacts of the C-terminal α5 helix of Gα_s_ with TM5 and the TM7/H8 hinge region of the receptor and leads to a more pronounced outward displacement of TM6 compared to the H_1_R-G_q_ complex. Furthermore, the ICL2 of H_2_R points deeper into the hydrophobic pocket formed between the αN, αN-β1 hinge and α5 of the Ras domain of G_s_ than ICL2 of H_1_R in the G_q_ complex, which is engaged in more polar contacts with the C-terminal end of the αN of G_q_. Moreover, since the H_2_R-G_s_ structure was obtained in lipid NDs, the revealed electrostatic interactions between basic residues of the αN of Gα_s_ and the polar headgroup of the lipids may indicate a role for the G protein anchoring to the membrane to facilitate receptor coupling, as described previously^35^.

In summary, the results show that our CF production pipeline allows to provide new structural insights into GPCRs with a good global resolution. It further opens up new avenues for the functional nanotransfer of membrane proteins from NDs to membranes of living cells^19–21^. Here, we used this approach to confirm the previously proposed homo-and heterooligomerization of the H_1_R and H_2_R using pull-downs and internalization assays. As the CF synthesized receptors are easily accessible for mutations or modifications, the nanotransfer strategy may also help to further characterize the GPCR interaction requirements and interfaces. We anticipate that the CF production of GPCRs will become more broadly applicable for structure determination of full-length and less engineered receptors in complex with heterotrimeric G proteins and in absence of detergents.

## Methods

### Lysate preparation

*E. coli* S30 lysates were prepared as described previously^52, 53^. Briefly, a pre-culture containing 150 mL LB media was inoculated with *E. coli* A19 cells and incubated at 37 °C with shaking (180 rpm) over night. A fermenter was filled with 10 L YPTG medium (16 g/L peptone, 10 g/L yeast, 5 g/L NaCl, 100 mM glucose, 22 mM KH_2_PO_4_, 40 mM K_2_HPO_4_). The fermenter was inoculated with 100 mL of the pre-culture and grown at 37 °C with vigorous stirring (300 rpm) and approx. 1 mL antifoam Y-30 emulsion (Sigma-Aldrich, Taufkirchen, Germany). When an OD_600_ of 3.5 to 4 was reached, the culture was rapidly cooled to 20 °C and subsequently harvested by centrifugation at 5,000 x g and 4 °C for 30 min. Cells were resuspended in 300 mL S30 buffer A (14 mM Mg(OAc)_2_, 60 mM KCl, 6 mM β-mercaptoethanol, 10 mM Tris-acetate, pH 8.2) and centrifuged at 10,000 x g and 4 °C for 10 min. This step was repeated twice and the last centrifugation was extended to 30 min. The washed pellet was resuspended in 110 % (w/v) S30 buffer B (14 mM Mg(OAc)_2_, 60 mM KCl, 1 mM DTT, 10 mM Tris-acetate, pH 8.2) and cells were disrupted using a French press with constant pressure of 20,000 psi. Disrupted cells were centrifuged twice at 30,000 x g and 4 °C for 30. The supernatant was adjusted to 400 mM NaCl and incubated at 42 °C for 45 min for a mRNA run-off. The solution was dialysed 2x at 4 °C against 5 L S30 buffer C (14 mM Mg(OAc)_2_, 60 mM KAc, 0.5 mM DTT, 10 mM Tris-acetate, pH 8.2) for 3 and 12 h, respectively. After a final centrifugation at 30,000 x g and 4 °C for 30 min, the supernatant was collected, aliquoted, flash frozen in liquid nitrogen and stored at −80 °C.

### Expression and purification of MSP1E3D1

MSP1E3D1 was expressed and purified as described previously^54^. For removal of the N-terminal His_6_-tag, the protein was TEV digested followed by a reverse IMAC. The MSP1E3D1 solution was set to 1 mM DTT before TEV was added in a MSP1E3D1 to TEV ratio of 1:25. The mixture was dialyzed against 1 mM DTT, 0.5 mM EDTA, 50 mM Tris-HCl, pH 8.0 at 4°C over night. Before loading on a pre-equilibrated HiTrap™ IMAC FF column (Cytiva, Munich, Germany), the mixture was centrifuged at 20,000 x g and 4 °C for 10 min. The flow through was collected and the column was washed with 10 column volumes (CVs) equilibration buffer (20 mM IMD, 100 mM NaCl, 20 mM Tris-HCl, pH 8.0). The flow through and wash fractions were concentrated to 3-5 mg/mL using Amicon ultrafiltrators (10 kDa MWCO, Merck Millipore, Darmstadt, Germany). All fractions were combined and dialyzed 2x over night at 4 °C against 5 L 10 % (v/v) glycerol, 300 mM NaCl, 40 mM Tris-HCl, pH 8.0. Until ND formation, the protein was flash frozen in liquid nitrogen and stored at −80 °C.

### Nanodisc formation

Purified MSP1E3D1 was incubated with the respective lipid and supplemented with 0.1 % DPC. For each lipid, NDs were assembled at defined MSP1E3D1 to lipid ratios (1:80 for DOPG, 1:80 for DOPC, 1:85 for DEPG, 1:110 for DMPG)^16, 54^. Solutions were incubated at RT for 1 h with gentle stirring, before being dialyzed 3x for at least 12 h at RT against 100 mM NaCl, 10 mM Tris-HCl, pH 8.0. The mixture was centrifuged at 30,000 x g and 4 °C for 20 min. NDs were concentrated to 500-1000 µM using Centriprep concentrating units (10 kDa MWCO, Merck Millipore, Darmstadt, Germany). Lipids and DPC were purchased from Avanti Polar Lipids (Alabaster, USA).

### CF expression

Coding sequences of H_1_R-Strep, H_2_R-Strep and FFAR_2_-Strep were synthesized from Twist Bioscience and cloned into vector pET29b. Constructs comprised the full-length GPCRs containing a N-terminal H-tag for improved expression^55^ as well as a C-terminal Strep-tag for affinity chromatography. For the nanotransfer or determination of expression levels in different NDs, a C-terminal mNG-fusion was introduced. CF protein expression was performed as described before^52, 56^. Optimal Mg^2+^ concentrations were determined for each DNA template by screening within a concentration range of 14 – 22 mM. Analytical scale reactions were carried out in 24 well plates using Mini-CECF reactors. Preparative scale reactions were carried out in 3 mL Slide-A-Lyzer cassettes (10 kDa MWCO, Merck Millipore, Darmstadt, Germany) in combination with custom-made FM containers^57^. The RM to FM ratios were 1:17. For cotranslational solubilization, all GPCRs were synthesized in presence of 60 µM NDs. Reactions were incubated at 30 °C for 16 to 20 h with gentle shaking. After expression, RMs were harvested and centrifuged at 18,000 x g and 4 °C for 10 min to remove precipitates. For purification of GPCR/ND complexes, RMs were diluted 1:3 in buffer A (100 mM NaCl, 20 mM HEPES, pH 7.4). Gravity flow columns containing StrepII-Tactin resin (IBA, Goettingen, Germany) were equilibrated in buffer A and samples were loaded and re-loaded twice. The columns were washed with 10 CVs buffer A and eluted in 4 to 5 CVs [buffer A + 25 mM d-desthiobiotin]. The samples were subsequently concentrated using Amicon Ultra - 0.5mL units (MWCO 50 kDa, Millipore, Merck, Darmstadt, Germany).

### Expression and purification of Nb35

Nb35 was expressed and purified as described previously^50^. Briefly, Nb35 was expressed in *Escherichia coli* BL21 cells. After lysis, it was purified using nickel affinity chromatography and finally subjected to size exclusion chromatography on a Superdex 200 10/300 gel filtration column (GE Healthcare) in 20 mM HEPES pH 7.5, 150 mM sodium chloride. Purified Nb35 was concentrated, flash frozen, and stored at –80°C until further use.

### Nanotransfer

Flag-H_1_R and Flag-H_2_R were ordered from Addgene (#66400 & #66401) and de-tangonized using standard site-directed mutagenesis^58^. Fluorescence microscopy and pulldowns with transferred GPCRs were performed as described before^19^. Briefly, for fluorescence microscopy, HEK293T cells were seeded at a density of 2 x 10^5^ cells/well onto glass coverslips in a 12-well plate. After 24h, fresh Dulbecco’s Modified Eagle Medium (DMEM) with 0.5 µM of Strep-purified GPCR/ND complexes was added and cells were incubated for 4h. Afterwards, they were washed 5 times with Dulbecco’s Phosphate Buffered Saline. To stimulate GPCR activation and subsequent internalization, the cells were incubated with DMEM containing 100 µM histamine. After 1h, cells were washed and fixed with RotiHistofix. For pulldowns, cells were seeded into 6-well plates and transfected with Flag-H_1_R and Flag-H_2_R. The next day, 0.5 µM GPCR/ND complexes were added for 16h. Cells were then lysed, and anti-Strep pulldowns were performed using magnetic Strep-Tactin beads.

### Size exclusion chromatography

SEC was carried out at 12 °C, using an Äkta purifier system (Cytiva, Munich, Germany) and Increase Superose 6 5/150 or 3.2/300 columns (Cytiva, Munich, Germany). The columns were equilibrated in sterile-filtrated, degased and pre-cooled buffer [100 mM NaCl, 20 mM HEPES, pH 7.4] before injecting the affinity purified protein samples. Flow rate was 0.15 mL/min for Superose 6 5/150 and 0.05 mL/min for Superose 6 3.2/300. UV absorbance was recorded at 280 nm and data were plotted using *GraphPad Prism*.

### Expression and purification of heterotrimeric G_s_

G_s_ heterotrimer was expressed and purified as previously described^59^. Briefly, *Trichoplusia ni* (*T. ni*) insect cells using baculoviruses generated by the BestBac method (Expression Systems) were used for expression of G_s_ heterotrimer. One baculovirus encoding the human Gα_s_ short splice variant and another separate baculovirus encoding both the Gβ_1_ and Gγ_2_ subunits, with a histidine tag and HRV 3C protease site inserted at the amino terminus of the β-subunit were used. *T. ni* cells were infected with the baculoviruses followed by incubation of 48 hours at 27 °C. After harvest by centrifugation, cells were lysed in [10 mM Tris, pH 7.5, 100 µM MgCl_2_, 5 M β-ME, 20 μM GDP and protease inhibitors]. The membrane fraction was collected by centrifugation and solubilized in [20 mM HEPES, pH 7.5, 100 mM NaCl, 1 % Na-cholate, 0.05 % DDM, 5 mM MgCl_2_, 5 mM β-ME, 5 mM IMD, 20 μM GDP and protease inhibitors]. After homogenization with a Dounce homogenizer, the solubilization reaction was incubated for 45 min at 4 °C. After centrifugation, the soluble fraction was loaded onto HisPur Ni-NTA resin (Thermo Scientific, Langenselbold, Germany) followed by a gradual detergent exchange into 0.1 % DDM. The protein was eluted in buffer supplemented with 200 mM IMD and dialyzed overnight against [20 mM HEPES, pH 7.5, 100 mM NaCl, 0.05 % DDM, 1 mM MgCl_2_, 5 mM β-ME and 20 μM GDP] together with HRV 3C protease to cleave off the amino-terminal His_6_-tag. Cleaved His_6_-tag, uncleaved fractions and 3C protease were removed by Ni-chelated Sepharose. The cleaved G protein was dephosphorylated by lambda protein phosphatase (NEB, Frankfurt, Germany), calf intestinal phosphatase (NEB, Frankfurt, Germany), and antarctic phosphatase (NEB, Frankfurt, Germany) in the presence of 1 mM MnCl_2_. Lipidated G_s_ heterotrimer was isolated using a MonoQ 5/50 GL column (Cytiva, Munich, Germany). The protein was bound to the column in buffer A [20 mM HEPES, pH 7.5, 50 mM NaCl, 1 mM MgCl_2_, 0.05 % DDM, 100 µM TCEP, 20 µM GDP] and washed in buffer A. The G_s_ heterotrimer was eluted with a linear gradient of 0 to 50 % buffer B [buffer A + 1 M NaCl]. The main peak containing isoprenylated G_s_ heterotrimer was collected and dialyzed against [20 mM HEPES, pH 7.5, 100 mM NaCl, 0.02 % DDM, 100 µM TCEP and 20 µM GDP]. The protein was concentrated to 250 µM, 20 % glycerol was added and the protein was flash frozen in liquid nitrogen and stored at –80 °C until use.

### Cryo-EM sample preparation

The H_2_R/ND/G_s_/Nb35-His complex was formed co-translationally. Reactions were supplemented with final concentrations of 15 ng/µL H_2_R-Strep template, 60 µM NDs(DOPG) without His-tag, 10 µM purified G_s_ heterotrimer and 15 µM Nb35-His. Reactions were incubated for 16 h at 30 °C with gentle shaking. Samples were harvested by centrifugation at 18,000 x g and 4 °C for 10 min and subsequently incubated with 1 U/µL apyrase on ice for 90 min. The H_2_R/ND/G_s_/Nb35-His complex was purified by IMAC at 4 °C. Samples were diluted 1:3 in IMAC buffer A [100 mM NaCl, 20 mM HEPES, pH 7.4] and loaded and reloaded twice on a a pre-equilibrated gravity-flow IMAC column. The column was washed with 4 column volumes IMAC buffer A and 4 column volumes IMAC buffer B (IMAC buffer A + 30 mM IMD). Samples were eluted using IMAC elution buffer (IMAC buffer A + 300 mM IMD) and concentrated using Amicon Ultra – 0.5 mL units (50 kDa MWCO, Merck Millipore, Darmstadt, Germany). SEC was performed as described above. Complex containing fractions were pooled and concentrated again using Amicon Ultra – 0.5 mL units (50 kDa MWCO, Merck Millipore, Darmstadt, Germany).

### Cryo-EM data acquisition

Two datasets were collected using the same settings, but different concentrations. Samples were prepared as previously described^60^. Specifically, the sample was concentrated to 1.4 mg/ml for the first and 2.8 mg/ml for the second dataset. C-flat grids (Protochips; CF-1.2/1.3-3Cu-50) were prepared by glow-discharging them with a PELCO easiGlow at 15 mA for a duration of 45 seconds. 3 µL of the sample were promptly applied to the grids and immediately plunge-frozen in liquid ethane with the use of a Vitrobot Mark IV (Thermo Fischer). This process was conducted at 4 °C with a relative humidity of 100%. Data was collected on a Glacios microscope (Thermo Fischer), operating at 200 kV and equipped with a Selectris energy filter (Thermo Fischer) with a slit with of 10 eV. Movies were recorded using a Falcon 4 direct electron detector (Thermo Fischer) at a nominal magnification of 130.000 which is equal to a calibrated pixel size of 0.924 Å per pixel. The dose rate for collection was set to 5.22 e^-^ per pixel with a total dose of 50 e^-^ per Å^2^. 46896 movies were automatically gathered using EPU software (v.2.9, Thermo Fischer) with a defocus range of −0.8 µm to −2.0 µm and stored in EER (electron-event representation) format.

### Cryo-EM image processing

The dataset was processed using cryoSPARC (v.4) (Extended Data Fig. S4). The preprocessing of the movies entailed patch-based motion correction, patch-based CTF estimation and filtering based on the CTF fit estimates using a cutoff at 5 Å. This resulted in a remaining dataset of 41594 micrographs.

Distinct 2D classes were identified and then utilized for further template-based particle picking. This process yielded a collection of 30.5 million particles. The particles were extracted within a box size of 288 pixels and then Fourier cropped to 72 pixels. Subsequent rounds of 2D classification were applied to refine the stack further, resulting in 1.5 M particles. These particles were then used for ab-initio 3D reconstruction. Particles from the top four reconstructions were merged and further refined by heterogeneous refinement, followed by another round of ab-initio 3D reconstruction. A final stack consisting of 425 thousand particles was utilized for a Non-Uniform refinement, resulting in a consensus map with a resolution of 3.5 Å. To enhance the quality of the map further, a local refinement was executed, resulting in a final map with a resolution of 3.4 Å.

### Model building and refinement

The preliminary atomic structure was computed using ModelAngelo (v.0.3). Subsequently, the generated structure was manually inspected in Coot (v.0.9) and iteratively refined using *phenix.real_space_refine* within Phenix (v.1.19). Further enhancement of the density map’s quality was accomplished through *phenix.density_modification*. Validation reports were generated by MolProbity. The final density map and corresponding atomic structure have been deposited in the Electron Microscopy Data Bank and the Protein Data Bank. These submissions can be found with the PDB ID 8POK and the EMDB ID 17793, respectively. All structural data was visualized using ChimeraX.

### Docking calculations

The reported H_2_R structure, as well as the H_1_R structure in an active conformation (PDB ID 7DFL), were prepared for docking calculations by adding protons and consequent energy minimization, acylation of the N-terminus, N-methylation of the C-terminus, and conversion of the mutated back to the wild-type or construction of the unmodeled residues using the Molecular Operating Environment (MOE, version 2022.02). The predicted binding modes of the compounds were energy-minimized using the MMFF94x force field. Binding pocket residues were relaxed through energy minimization in presence of the different compounds using the AMBER force field, as implemented in MOE. The following softwares were used for docking calculations: Autodock Vina^61^, DOCK3.7^62^, the OpenEye programs FRED and HYBRID, and SEED^63^. UCSF Chimera^64^ was used for generating the pdbqt format file used in Autodock Vina. The top-ranked binding modes from each docking calculation were inspected visually and those that had unfavorable interactions not considered further. RDKit software was used for RMSD calculations (RDKit: Open-source cheminformatics. https://www.rdkit.org). Pymol was used for visualization and pose evaluation.

### Cell culture & transient transfection for surface ELISA and BRET assays

Wildtype and mutant H_1/2_R and the G protein biosensors, G_q_-CASE and G_s_-CASE (plasmids are available from Addgene (https://www.addgene.org/browse/article/28216239/) (Reference to PMID: 34516756), were transiently expressed in HEK293A cells grown in Dulbecco’s Modified Eagle’s Medium (DMEM) supplemented with 2 mM glutamine, 10% fetal calf serum, 0.1 mg/mL streptomycin, and 100 units/mL penicillin at 37 °C with 5% CO_2_. Each mL of resuspended cells (300,000 cells / mL) was mixed with 500 ng receptor, 500 ng G-CASE and 3 mL PEI solution (1 mg/mL) (Merck KGaA, Darmstadt, Germany). 100 µL transfected cells were seeded per well onto 96-well plates (Brand, Wertheim, Germany) and grown for 48 hours at 37 °C with 5% CO_2_. White plates were used for BRET experiments and transparent, flat bottom 96-well plates (Brand, Wertheim, Germany) were used for the assessment of receptor surface levels. Absence of mycoplasma contamination was routinely confirmed by PCR.

### Assessment of receptor surface expression through live-cell ELISA

To quantify cell surface receptor expression, HEK293A cells transfected with G-CASE and pcDNA or N-terminally FLAG-tagged H_1/2_R constructs were grown for 48 hours in transparent 96-well plates (Brand, Wertheim, Germany) and washed once with 0.5% BSA (Merck KGaA, Darmstadt, Germany) in PBS. Next, cells were incubated with a rabbit anti-FLAG M2 antibody (142 ng/mL) (Cell Signaling Technology, Danvers, MA, USA) in 1% BSA–PBS for 1 h at 4 °C. Following incubation, the cells were washed three times with 0.5% BSA–PBS and incubated with a horseradish peroxidase-conjugated goat anti-rabbit antibody (30 ng/ml) (Cell Signaling Technology, Danvers, MA, USA) in 1% BSA–PBS for 1 h at 4 °C. The cells were washed three times with 0.5% BSA/PBS, and 50 µl of the 3, 3’, 5, 5’ tetramethyl benzidine (TMB) substrate (BioLegend, San Diego, CA, USA) was added. Subsequently, the cells were incubated for 30 min and 50 µl of 2 M HCl was added. The absorbance was read at 450 nm using a BMG ClarioStar Plus plate reader.

### BRET-based G protein activation experiments

G protein activation experiments with the G-CASE biosensors were conducted as previously described (Reference to PMID: 34516756). Briefly, transfected cells grown for 48 hours in 96-well plates were washed with Hanks′ Balanced Salt solution (HBSS) and incubated with 1/1,000 dilution of furimazine stock solution (Promega, WI, USA). After incubation for 2 minutes, BRET was measured in three consecutive reads followed by addition of agonist (histamine/amthamine) solutions or vehicle control and subsequent BRET reads. All experiments were conducted at 37 °C. Nluc emission intensity was selected using a 470/80 nm monochromator and cpVenus emission using a 530/30 nm monochromator in a CLARIOstar plate reader with an integration time of 0.3 seconds.

### BRET data analysis

BRET ratios were defined as acceptor emission/donor emission. The basal BRET ratio before ligand stimulation (BRET_basal_) was defined as the average of at least three consecutive reads. To quantify ligand-induced changes, D BRET was calculated for each well as a percent over basal [(BRET_stim_− BRET_basal_)/BRET_basal_] × 100). Next, the average DBRET of vehicle control was subtracted. Data were analysed using Prism 9.5 software (GraphPad, San Diego, CA, USA). Data from BRET concentration–response experiments were fitted using a three-parameter fit. Data from cell surface ELISA experiments were corrected for background by subtracting the values obtained for pcDNA-transfected cells.

## Acknowledgements

We thank Birgit Schäfer for helpful discussions and technical assistance. We are further grateful to Ulrich Ermler for valuable help in structure calculation, to Betsy White for help with G protein expression and purifications and to Juliane Bernhard for artwork. The work was funded by the DFG project BE1911/9-1, by the Center for Biomolecular Magnetic Resonance and by the LOEWE project GLUE of the state of Hessen. LOEWE GLUE financed the theses of ZK, SU and MP. Financial support was further obtained by the Fonds der Chemischen Industrie (SK-208/16).

## Author contributions

Sample preparations and biochemical studies were done by ZK and DH. KS, DJ, KP and AM performed cryo-EM data acquisition and analysis. DH provided heteromeric G-proteins. SU performed the nanotransfer and analysis in HEK293T cells. HS, DH and SP characterized mutants and analyzed ligand selectivity. MP and PK performed molecular docking studies. VD provided essential support and infrastructure. FB and DH conceived the project. All authors contributed to manuscript writing, data analysis, reading and approving the final version of the manuscript.

## Extended data

**Fig. S1.**
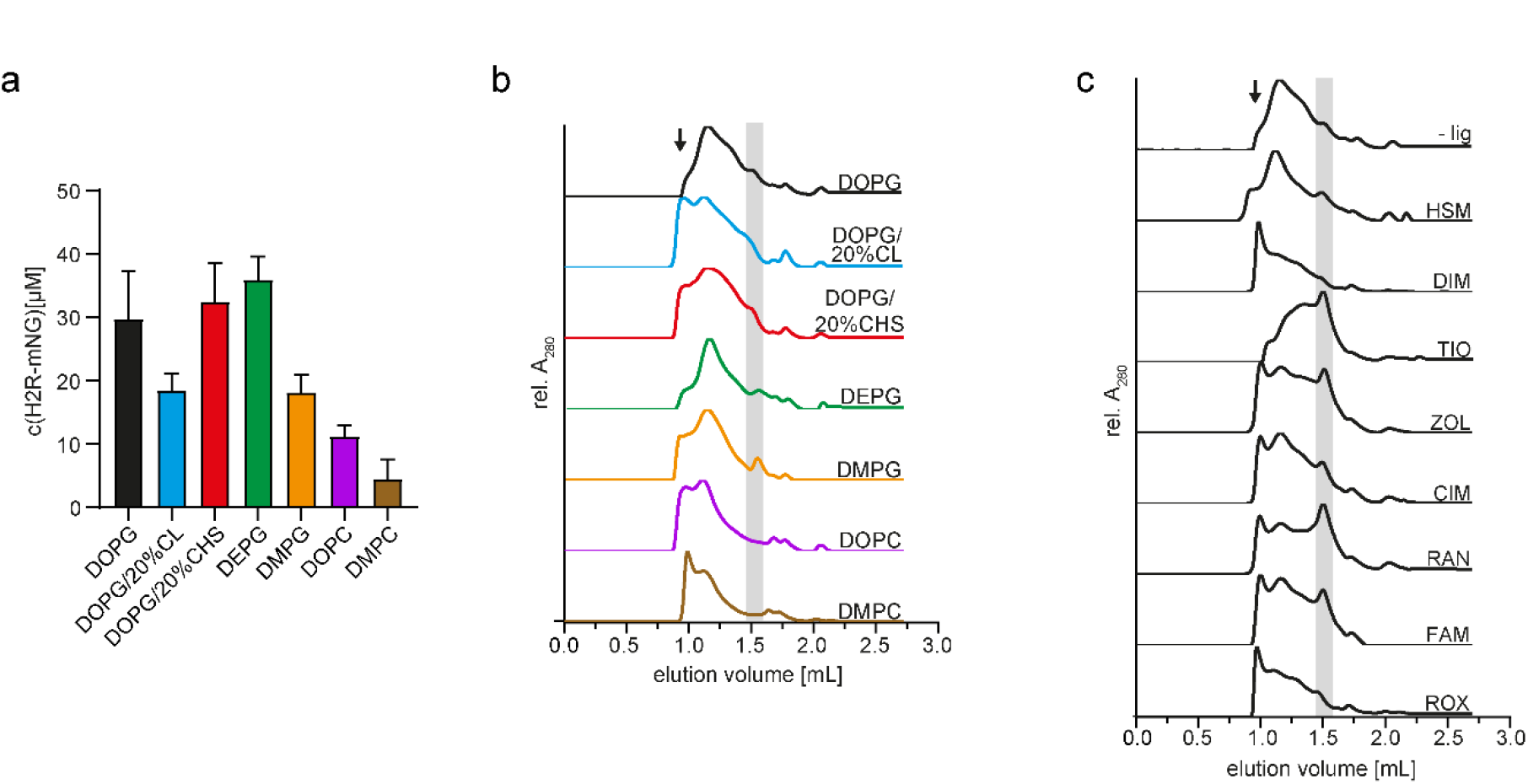
Effect of lipids and supplied ligands on H_2_R solubilization and sample quality. **a.** Cotranslational solubilization of H_2_R-mNG in different lipid environments. H_2_R-mNG was synthesized in presence of 60 µM NDs pre-assembled with the indicated lipids. The H_2_R-mNG/ND complexes were Strep-purified and quantified according to the mNG fluorescence. Data are presented as mean s.d. (n = 3 individual experiments). **b.** SEC profiles of Strep-purified H_2_R/ND complexes CF synthesized in presence of 60 µM preformed NDs containing membranes with the indicated lipids. **c.** SEC profiles of H_2_R CF synthesized in presence of 60 µM NDs (DOPG) and in presence of supplied ligands. –lig: without ligand; HSM: 5 mM histamine; DIM: 5 mM dimaprit; Tio: 200µM tiotidine; ZOL: 200µM zolantidine; CIM: 500 µM cimetidine; RAN: 200 µM ranitidine; FAM: 200 µM famotidine; ROX: 200 µM roxatidine. SEC was performed with a Superose 6 3.2/300 column at a flow rate of 0.05 mL/min. V, void volume; Fractions of proposed folded H_2_R are indicated in grey^13^.

**Fig. S2.**
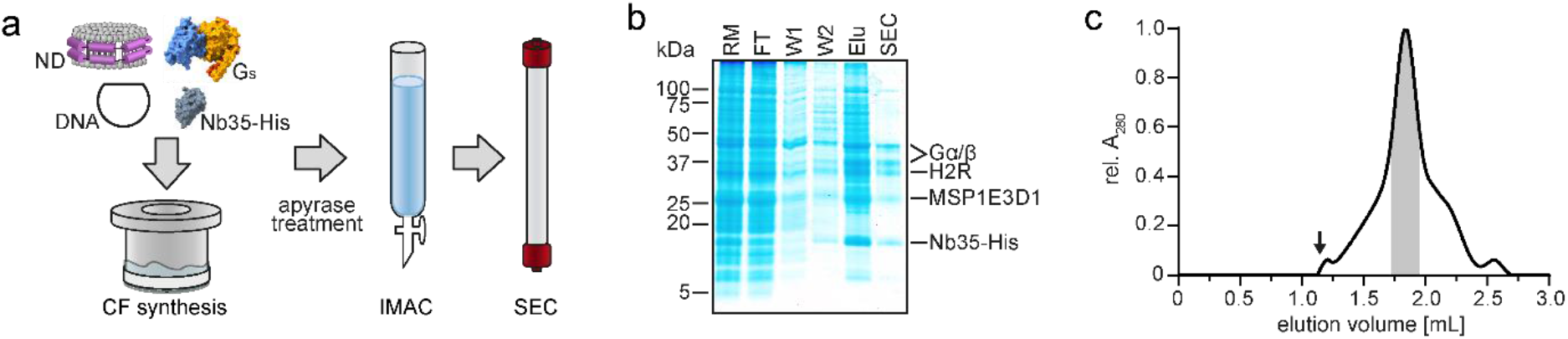
Cryo-EM sample production workflow. **a.** H_2_R is synthesized in presence of empty, pre-formed NDs, G_s_ heterotrimer and Nb35-His. After apyrase treatment, the sample is subsequently purified by IMAC and SEC. **b.** SDS-PAGE of the H_2_R/ND/G_s_/Nb35-His complex purification process. RM: reaction mix, FT: flow through, W1: first wash, W2: second wash, Elu: elution. **c.** SEC profile of the purified H_2_R/ND/G_s_/Nb35-His complex. Gelfiltration was performed with a Superose 6 5/150 Increase column at a flow rate of 0.15 mL/min. The void volume is indicated by an arrow. The fractions collected and concentrated for cryo-EM analysis are indicated in grey.

**Fig. S3.**
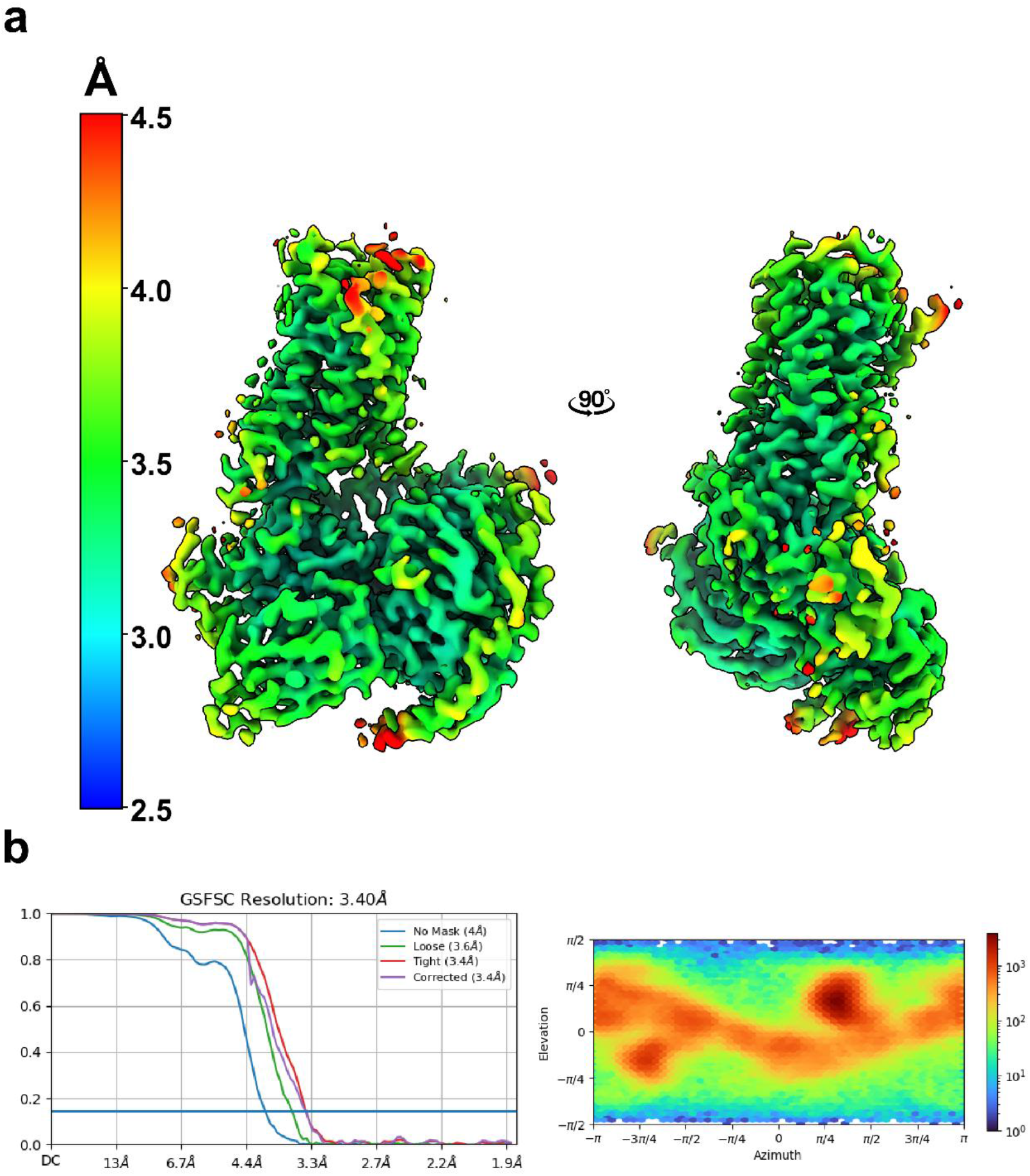
Resolution of the H_2_R/G_q_ complex. **a.** Local resolution map of the H_2_R/G_q_ complex colored according to resolution. **b.** FSC curve and particle distribution for the displayed map.

**Fig. S4.**
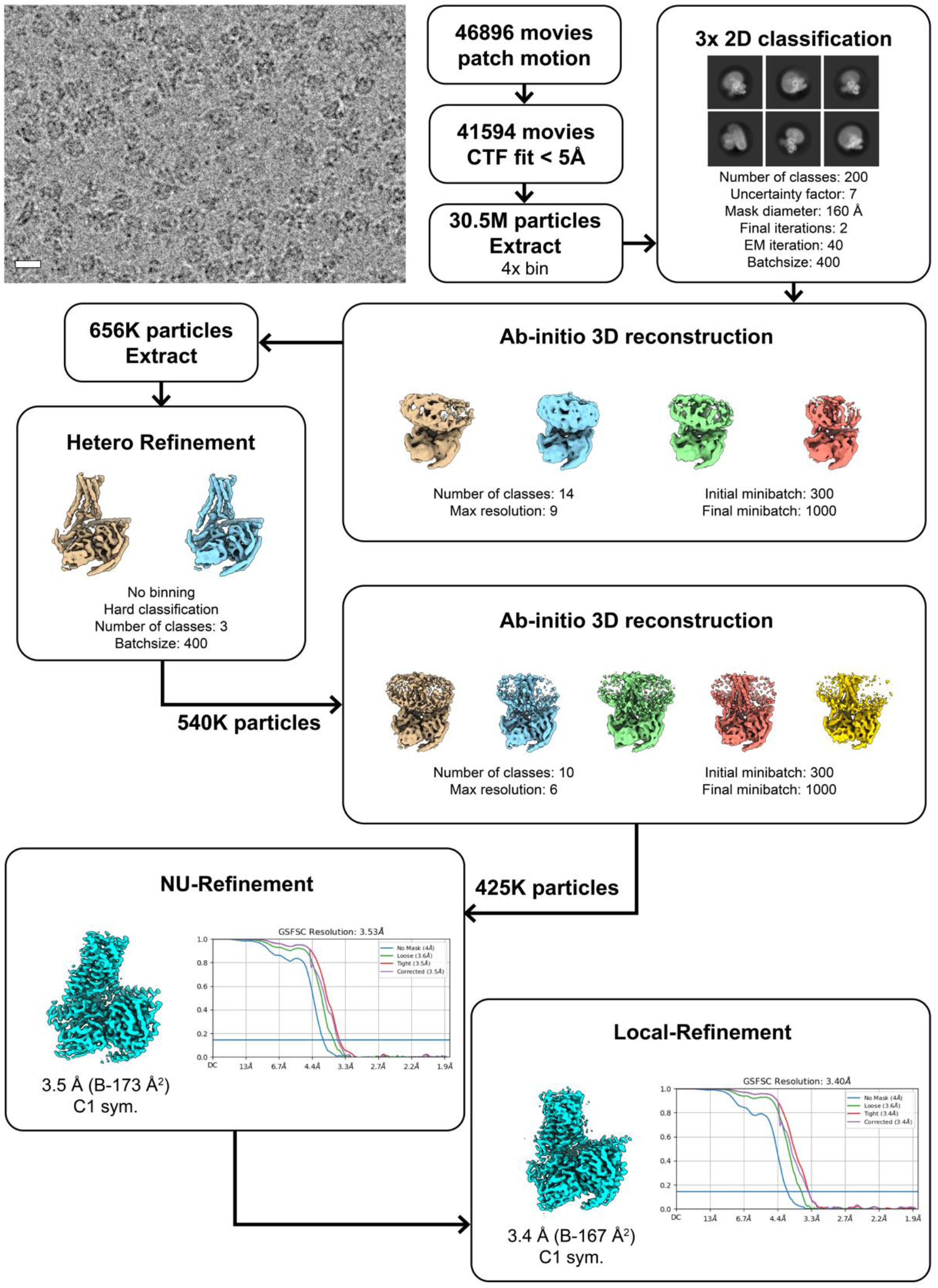
Cryo-EM data processing workflow. Processing flow chart for the H_2_R/G_q_ complex. Processing was performed in cryoSPARC. A representative cryo-EM micrograph, along with 2D-class averages and with a 100 Å scale bar in the micrograph is shown. Only the selected classes used for further processing were displayed in *ab-initio* 3D reconstruction and hetero refinement charts.

**Fig. S5.**
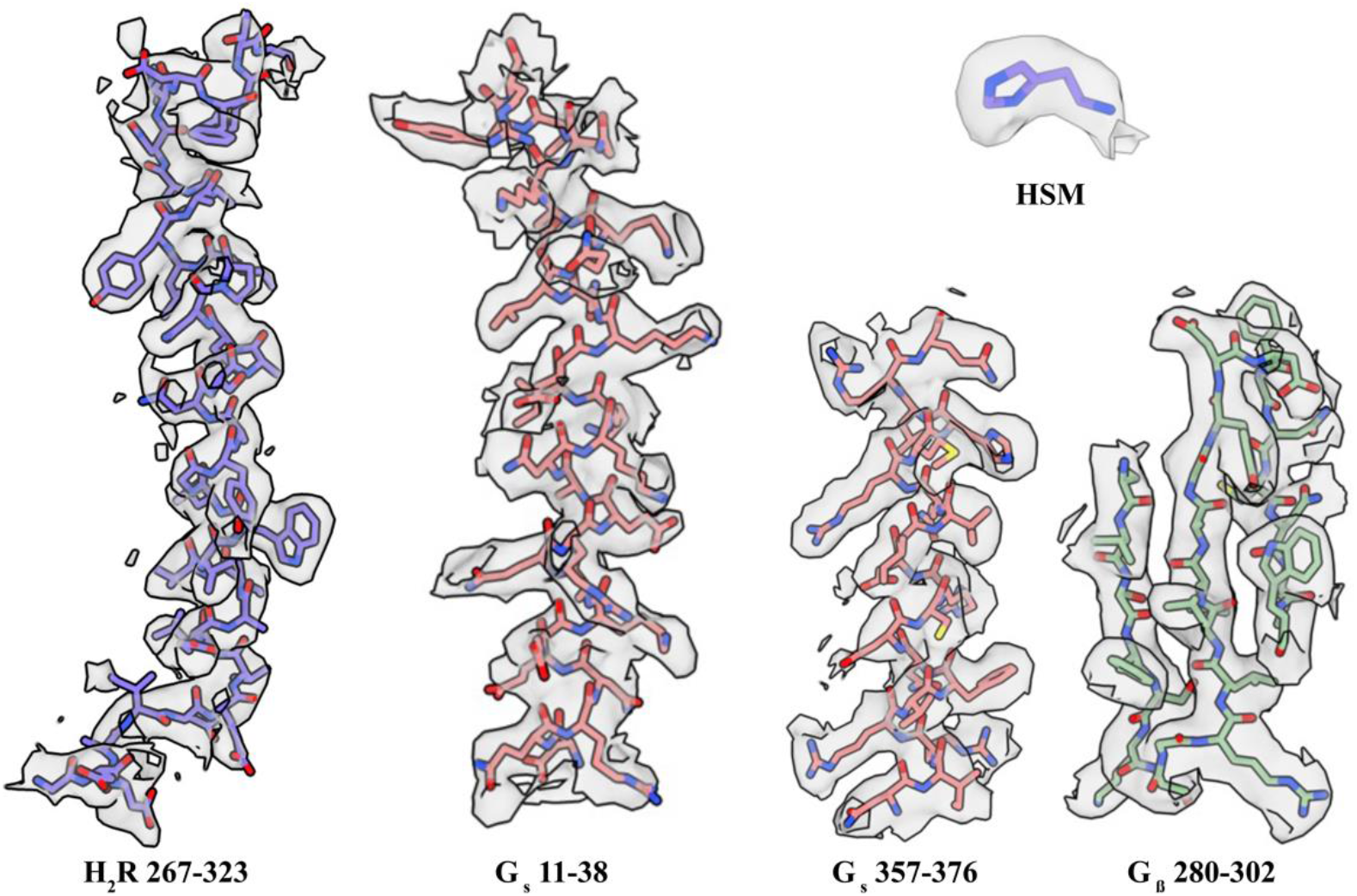
Representative cryo-EM densities of H_2_R with fitted models. Shown are the densities of TM7 from amino acid position 267-323 of H_2_R, of the α-helices 11-38 and 357-376 of G_s_, of the β-sheet 280-302 of G_β_ and of the agonist histamine (HSM).

**Fig. S6.**
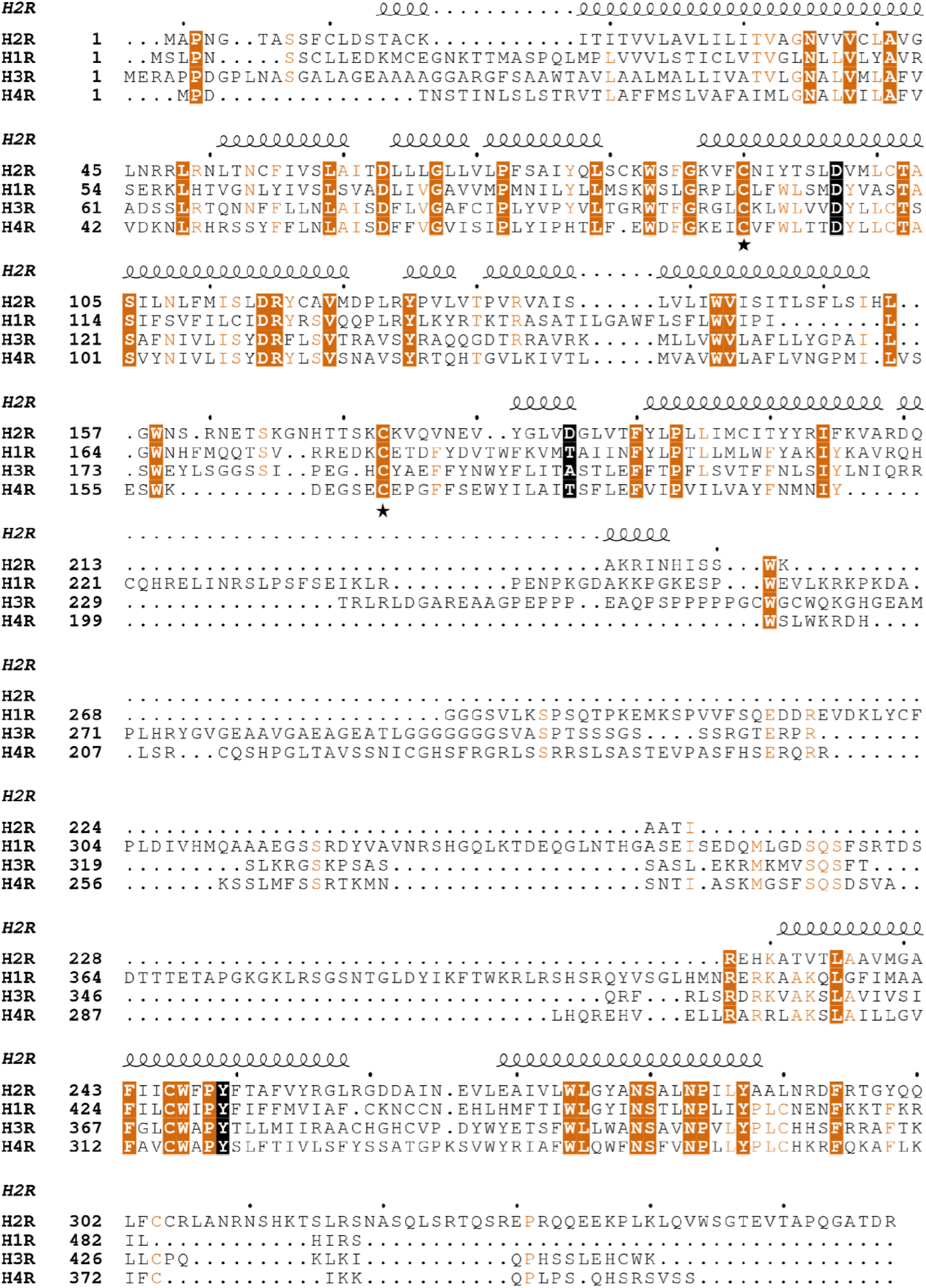
Amino acid sequence alignment of GPCR histamine receptors. The secondary structure symbols are illustrated according to the H_2_R structure. Conserved residues are highlighted. The coordinating residues of the ligand are marked with the corresponding boxes in black. Disulfide cysteines marked with asterisks. The alignment figure was made using MAFFT multiple sequence alignment program and ESPript.

**Fig. S7.**
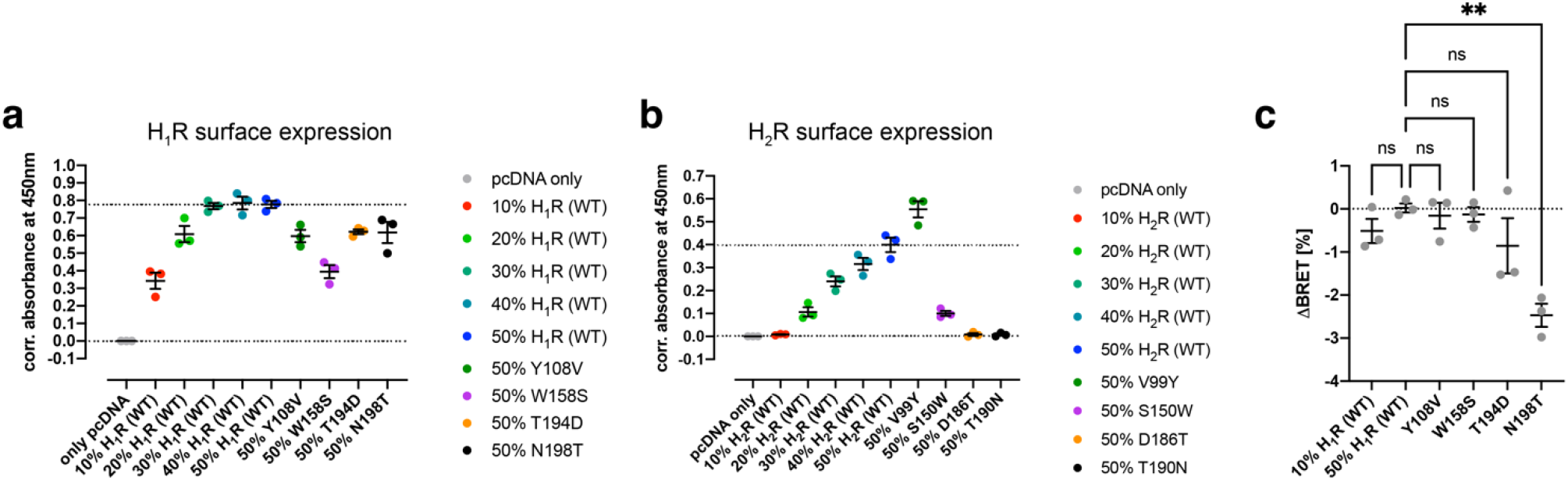
Comparative surface-expression levels of H_1_R and H_2_R constructs and analysis of the ΔBRET response in the H_1_R-dependent Gq activation assay in the presence of 100 uM amthamine. (**a, b**) The surface-expression levels of wild-type (WT) and H_1/2_R mutants were quantified using an ELISA against the N-terminal Flag tag. Different amounts of DNA encoding for the WT receptors was used for the transfection into HEK293A cells to adjust the surface-expression level of the WT to the receptor mutants to ensure comparable protein amounts in the plasma membrane for the BRET-based G protein signaling assays. Data obtained from three independent experiments. (**c**) Comparison of the ΔBRET response of the H_1_R (WT) and mutants in the presence of 100 uM amthamine showing a statistically significant decrease in the BRET signal of the H_1_R-mutant N198T. Statistical significance in (c) was tested using One-Way ANOVA followed by Fisher’s LSD test for multiple comparison.

**Fig. S8.**
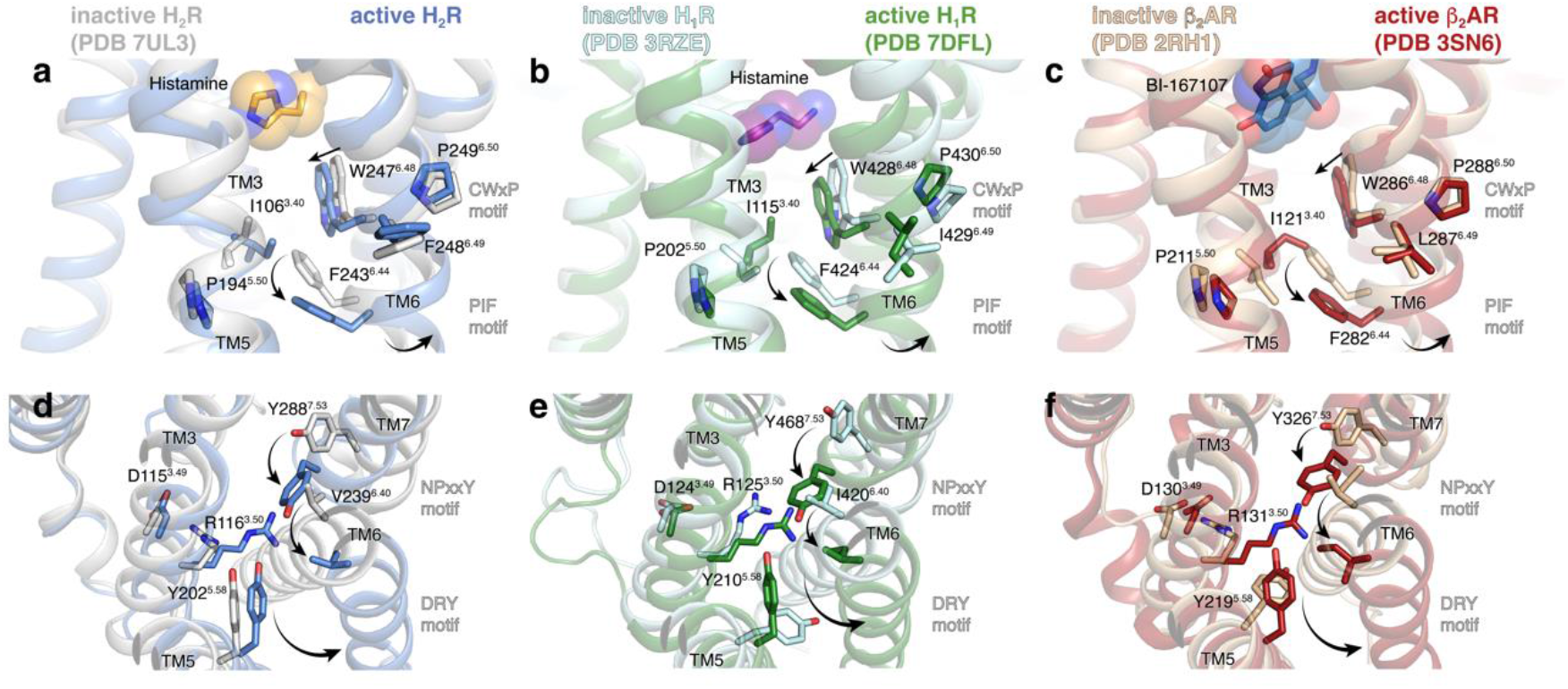
Comparison of the activation mechanism of the H_2_R, H_1_R and β_2_AR. **a.-f.** Close-up views of the conserved microswitch sequence motifs in the inactive and active structures of the (**a, d**) H_2_R, (**b, e**) H_1_R, and (**c, f**) β_2_AR. The receptors undergo similar conformational changes in the conserved (**a-c**) CWxP and PIF motifs and (**d-f**) NPxxY and DRY motifs, suggesting that these aminergic GPCRs share a similar overall allosteric activation mechanism between the ligand-binding site and the intracellular transducer-coupling cavity. The residues of the conserved microswitches are shown as stick models labeled by residue number and the corresponding Ballesteros-Weinstein numbering (superscript). The receptor-bound agonists are represented as spheres. Arrows indicate the direction of conformational changes upon activation of the receptors.

**Fig. S9.**
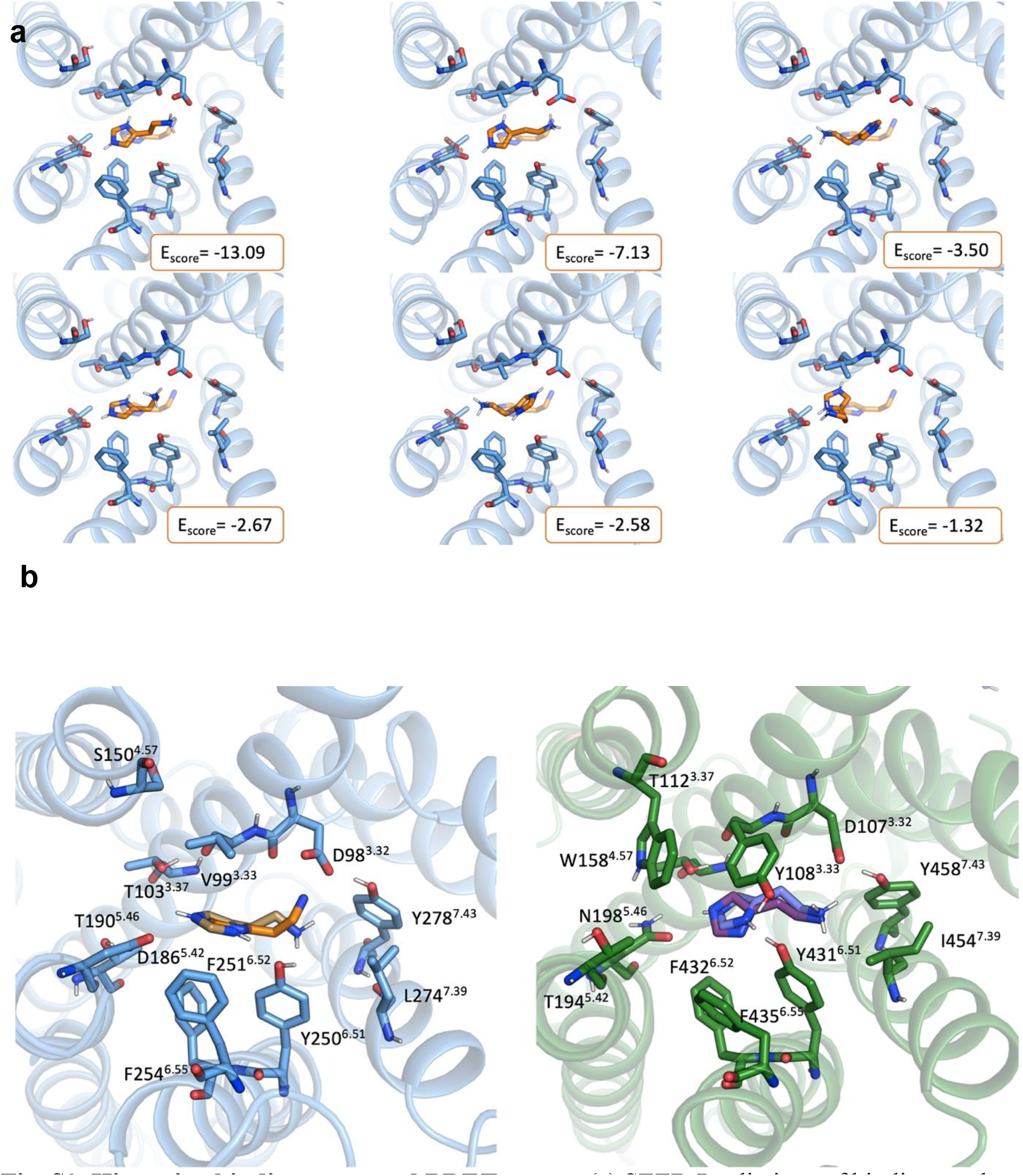
Histamine binding poses and BRET assays. (a) SEED Predictions of binding modes for histamine in the H_2_R. In the orange-framed boxes, the SEED energy score values are reported. The top-2-scored binding modes are reproducing the experimentally resolved orientation of the compound. (b) Left: predicted (sand) and experimentally resolved (orange) binding mode of histamine in the H_2_R; Right: predicted (violet) and experimentally resolved (red-purple) binding modes of histamine in the H_1_R. Binding mode predictions were obtained with AutoDock Vina.

**Fig. S10.**
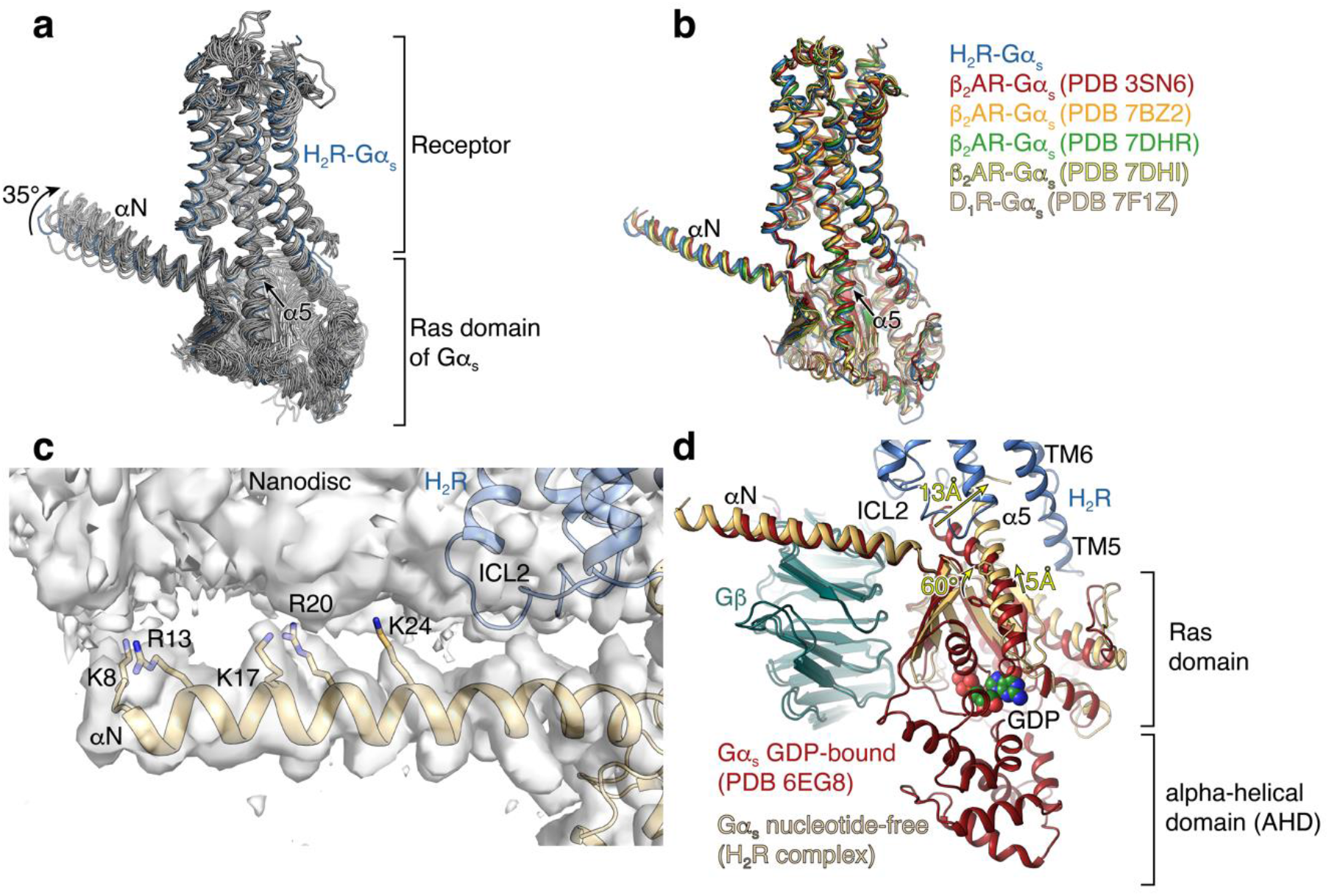
Orientation and interaction of the G protein G_s_ relative to the receptor and the nanodiscs lipid bilayer. a. Comparison of the G protein orientation in aminergic-receptor/G_s_ complexes (PDB IDs: 7XT9, 7XT8, 7XTB, 7XTC, 8DCR, 8DCS, 7X2F, 7F10, 7X2D, 7F0T, 7F24, 7F1Z, 7F23, 7XJI, 7XJH, 7S0G, 7DH5, 7JOZ, 7CKZ, 7LJC, 7CKW, 7CKY, 7CRH, 7LJD, 7JV5, 7JVQ, 7JVP, 7DHI, 7DHR, 7JJO, 7BZ2, 6NI3, 3SN6) showing differences in the relative orientation of the Gα_s_ subunit with respect to the receptor. The H_2_R-G_s_ complex is highlighted in blue. **b.** Complex structures showing similar receptor-G protein orientations in comparison to the H_2_R-Gα_s_ complex. **c.** Interaction of the αN helix of Gα_s_ (wheat) with the nanodisc membrane. Basic lysine and arginine residues that putatively form electrostatic interactions with the polar headgroups of the phospholipids are highlighted as stick models. The H_2_R is highlighted in blue. **d.** Structural changes in G_s_ upon coupling to the H_2_R. Major differences between the GDP-bound G_s_ structure (PDB ID 6EG8) and the nucleotide-free H_2_R-Gα_s_ complex involves the rotational translation of the α5 helix into the receptor core and the opening of the alpha-helical domain (AHD) to allow dissociation of the bound GDP (green spheres). Arrows indicate the direction of conformational changes upon activation of the receptors.

**Table S1:** *Cryo-EM data collection, refinement, and validation statistics*.

|  | H<sub>2</sub>R<br>(EMD- 17793)<br>(PDB 8POK) |
| --- | --- |
| **Data collection** |  |
| Microscope | Glacios |
| Energy filter and camera | Falcon 4<br>Selectris |
| Energy filter slit width | 10 eV |
| Voltage (kV) | 200 |
| Nominal magnification | 130,000 |
| Pixel size (Å) | 0.924 |
| Electron exposure (e<sup>-</sup>/Å<sup>2</sup>) | 50 |
| Total exposure time (s) | 8.17 |
| Number of frames per image | 50 |
| Total number of images | 46896 |
| Defocus range (μm) | - 0.8 to - 2.0 |
| **Image processing** |  |
| Processing software | cryoSPARC |
| Motion correction software | cryoSPARC |
| CTF estimation software | cryoSPARC |
| Particle selection software | cryoSPARC |
| Initial particle images (no.) | 30.5 M |
| Final particle images (no.) | 425 K |
| Final refinement software | cryoSPARC |
| Symmetry imposed | C1 |
| Map resolution (Å) | 3.4 |
| B-factor (Å<sup>2</sup>) | 119.3 |
| **Refinement statistics** |  |
| Initial model (PDB code) | 8POK |
| Modeling software | Coot, PHENIX,<br>model-angelo |
| **Model composition** |  |
| Non-hydrogen atoms | 8042 |
| Protein residues | 1024 |
| Water molecules | 0 |
| Ligands | 1 |
| **Mean B factors (Å<sup>2</sup>)** |  |
| Protein | 83.89 |
| Water molecules | - |
| Ligand | 68.59 |
| **R.m.s. deviations** |  |
| Bond lengths (Å) | 0.007 |
| Bond angles (°) | 0.978 |
| **Validation** |  |
| MolProbity score | 1.97 |
| Clashscore | 7.35 |
| Poor rotamers (%) | 0.00 |
| **Ramachandran plot** |  |
| Favored (%) | 89.29 |
| Allowed (%) | 10.71 |
| Disallowed (%) | 0.00 |

**Table S2:** AutoDock Vina docking scores for H2R selective agonists.

| VINA | H<sub>1</sub>R score | H<sub>2</sub>R score | VINA | H<sub>1</sub>R score | H<sub>2</sub>R score |
| --- | --- | --- | --- | --- | --- |
| Histamine | -4.8 | -4.1 | Dimaprit | -4.3 | -4.6 |
|  | -4.8 | -4.1 |  | -4.1 | -4.3 |
|  | -4.4 | -4.0 |  | -4.0 | -4.3 |
|  | -4.0 | -4.0 |  | -3.8 | -4.1 |
|  | -3.8 | -3.9 |  | -3.8 | -4.1 |
|  | -3.7 | -3.9 |  | -3.7 | -4.1 |
|  | -3.7 | -3.9 |  | -3.7 | -4.1 |
|  | -3.7 | -3.9 |  | -3.7 | -4.0 |
|  | -3.6 | -3.9 |  | -3.6 | -3.8 |
|  | -3.6 | -3.7 |  | -3.5 | -3.6 |
| Amthamine | -4.3 | -4.9 | Bis157 | -5.6 | -9.0 |
|  | -3.9 | -4.9 |  | -4.8 | -8.5 |
|  | -3.9 | -4.6 |  | -4.1 | -8.3 |
|  | -3.8 | -4.5 |  | -4.1 | -8.2 |
|  | -3.8 | -4.5 |  | -3.9 | -7.6 |
|  | -3.6 | -4.4 |  | -3.9 | -7.6 |
|  | -3.5 | -4.4 |  | -3.8 | -7.5 |
|  | -3.5 | -4.4 |  | -3.8 | -7.4 |
|  | -3.4 | -4.2 |  | -3.8 | -7.3 |
|  | -3.2 | -4.2 |  | -3.1 | -7.1 |

**Table S3.** Docking scores from different docking softwares for H1R/H2R-selective compounds.

|  | **FRED** | **HYBRID** | **DOCK** |
| --- | --- | --- | --- |
| Histamine | -9.6 | -6.9 | -41.6 |
| Amthamine | -8.2 | -6.2 | -39.3 |
| Dimaprit | -10.8 | -6.8 | nd |
| Bis-157 | nd | nd | -31.9 |

